# Transcription organizes euchromatin similar to an active microemulsion

**DOI:** 10.1101/234112

**Authors:** Lennart Hilbert, Yuko Sato, Hiroshi Kimura, Frank Jülicher, Alf Honigmann, Vasily Zaburdaev, Nadine L. Vastenhouw

## Abstract

Chromatin is organized into heterochromatin, which is transcriptionally inactive, and euchromatin, which can switch between transcriptionally active and inactive states. This switch in euchromatin activity is accompanied by changes in its spatial distribution. How euchromatin rearrangements are established is unknown. Here we use super-resolution and live-cell microscopy to show that transcriptionally inactive euchromatin moves away from transcriptionally active euchromatin. This movement is driven by the formation of RNA-enriched microenvironments that exclude inactive euchromatin. Using theory, we show that the segregation into RNA-enriched microenvironments and euchromatin domains can be considered an active microemulsion. The tethering of transcripts to chromatin via RNA polymerase II forms effective amphiphiles that intersperse the two segregated phases. Taken together with previous experiments, our data suggest that chromatin is organized in the following way: heterochromatin segregates from euchromatin by phase separation, while transcription organizes euchromatin similar to an active microemulsion.

---

In eukaryotes, DNA is packed inside the cell nucleus in the form of chromatin, which consists of DNA, proteins such as histones, and RNA. During the interphase of the cell cycle, chromatin exhibits a dynamic, three-dimensional (3D) organization (1,2) which is required for development and health (3–6). A prominent feature of this organization is the segregation into chromatin domains and interchromatin space (2). Chromatin domains are subdivided into mutually exclusive compartments, which contain either transcriptionally repressed heterochromatin (B compartment), or transcriptionally permissive euchromatin (A compartment) (7,8). Actually transcribed parts of euchromatin are unfolded and reach into the interchromatin space, which otherwise contains little chromatin (9–13). These aspects of 3D organization are conserved across many cell types (2). The segregation of heterochromatin from euchromatin has recently been suggested to be governed by a physical principle, phase separation (14,15). For euchromatin, however, a general principle that can explain its organization based on transcriptional activity, has not yet been proposed.

To determine the role of transcription in euchromatin organization, we used late blastula zebrafish cells. These cells do not display heterochromatin (SI Figure 1) or nucleoli (16,17) allowing us to focus on euchromatin. Furthermore, these cells divide approximately once per hour, which facilitates the frequent observation of transcription onset after mitosis and the concurrent establishment of euchromatin organization. We developed a protocol for simultaneous super-resolution imaging of DNA, RNA, and transcriptional activity within nuclei of intact cells by three-color STED microscopy (SI Figure 2). The DNA intensity profile inside nuclei was relatively smooth before transcription onset (SI Figure 3A-F), while a pattern of distinct DNA domains and DNA-depleted regions was present after transcription onset (Figure 1A and SI Figure 3G-I). Indeed, image contrast, a measure that quantifies how strongly the intensity in different areas of the nucleus differs (see Supplementary Methods), is significantly increased after transcription onset, reflecting the observed formation of DNA domains (Figure 1B and SI Figure 4A,B). To test whether the observed changes require transcription, we inhibited RNA polymerase II activity by injecting α-amanitin into zebrafish embryos. In this case, the formation of DNA domains did not occur (SI Figure 5). These results indicate that RNA polymerase II mediated transcription establishes a pattern of DNA domains.

**Figure 1.**
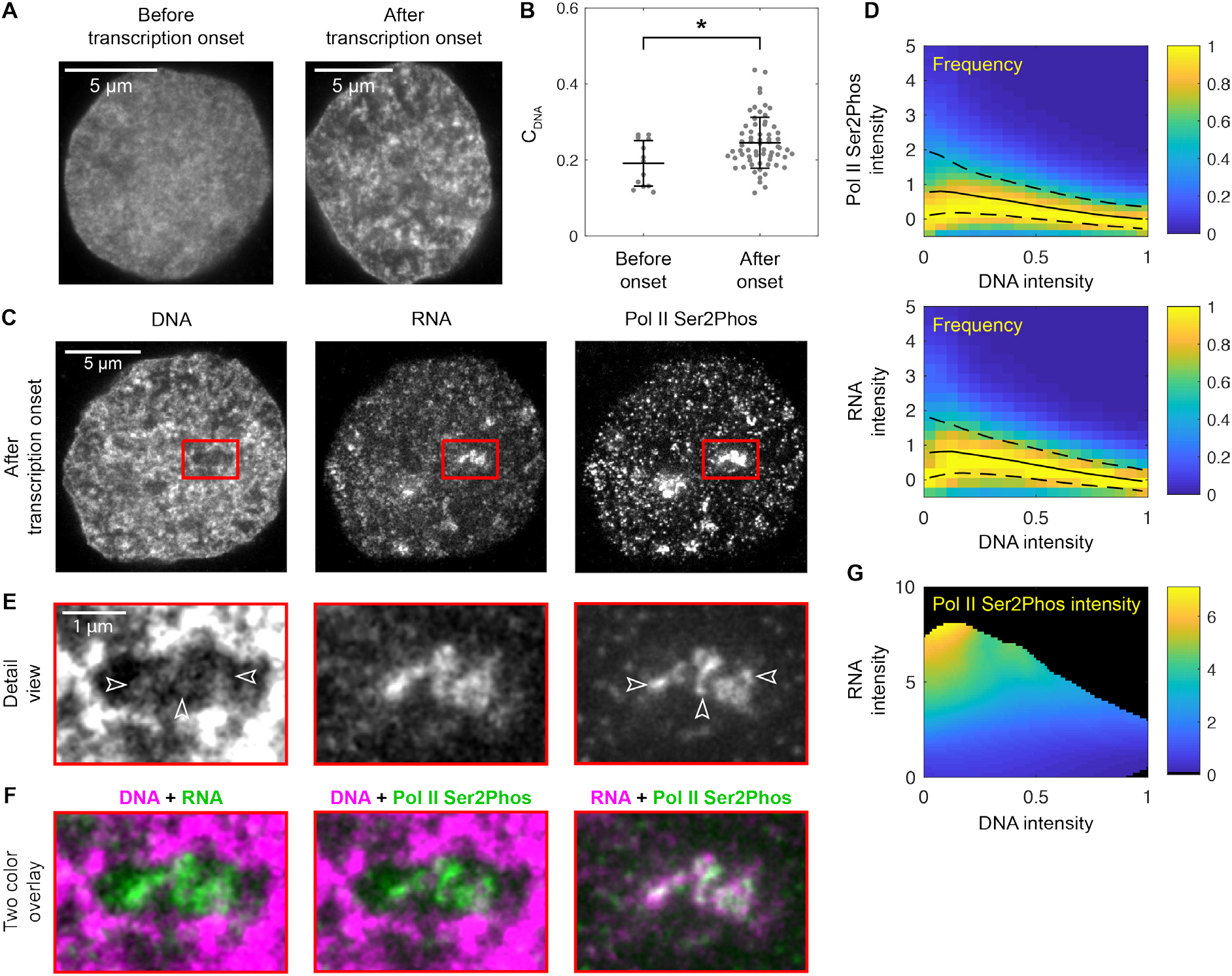
Transcription onset after mitosis establishes microenvironments that organize euchromatin. A) Representative STED super-resolution micrographs showing DNA intensity profiles in nuclear mid-sections before and after transcription onset. B) Image contrast (C_DNA_)in DNA intensity profiles from nuclear mid-sections before and after transcription onset (mean±std.dev., *p<0.05, permutation test, n=13, 66). C) Representative three-color STED micrographs showing DNA, RNA, and transcriptional activity (Pol II Ser2Phos) in a nuclear mid-section after transcription onset. D) Pol II Ser2Phos and RNA intensity distributions resolved by DNA intensity, intensity median (solid line) and quartile range (dashed lines) indicated. E, F) Zoomed view of a microenvironment, as indicated in panel C. G) Pol II Ser2Phos intensity resolved by DNA as well as RNA intensity.

The role of transcription in the establishment of a pattern of DNA domains might result from accumulation of the product of transcription, RNA, as well as from transcriptional activity itself. To dissect the individual contributions of RNA and transcriptional activity, we first investigated how they are spatially related to DNA domains in nuclei of transcriptionally active cells. As had been observed before (18) RNA and transcriptional activity (visualized by the Ser2-phosphorylated, elongating form of RNA polymerase II) were localized in regions that are generally depleted of DNA (Figure 1C). Quantitative analysis confirmed that the highest intensities of RNA and transcriptional activity occur in regions of low DNA intensity (Figure 1D). Inside these regions, low intensity DNA protrusions were retained (Figure 1E, arrowheads). It was specifically on these protrusions that we found peaks of transcriptional activity (Figure 1F), suggesting that the DNA protrusions represent transcribed DNA. In support of this interpretation, a two-dimensional analysis that resolved transcriptional activity by DNA and RNA intensity revealed that the highest intensity of transcriptional activity is consistently found in locations with low DNA intensity and maximal RNA intensity (Figure 1G). Together, our results suggest that transcription results in distinct RNA-enriched regions, or microenvironments. Transcribed DNA protrudes into these microenvironments while nontranscribed DNA forms domains that are spatially segregated from the RNA-enriched microenvironments.

The spatial segregation of RNA from chromatin suggests that accumulation of RNA in the nucleus might be required for the confinement of chromatin into distinct domains. To test this hypothesis, we inhibited transcription with flavopiridol. After flavopiridol treatment, nuclei retained a range of RNA levels, due to differences in the amount of nuclear RNA at the time flavopiridol was applied (SI Figure 4C). We could therefore determine how different amounts of RNA in the cell nucleus contribute to euchromatin organization. We found that DNA image contrast increased with greater amounts of RNA in the nuclei of flavopiridol-treated cells (Figure 2A). This suggests that RNA that accumulates in the nucleus establishes a pattern of chromatin domains.

**Figure 2.**
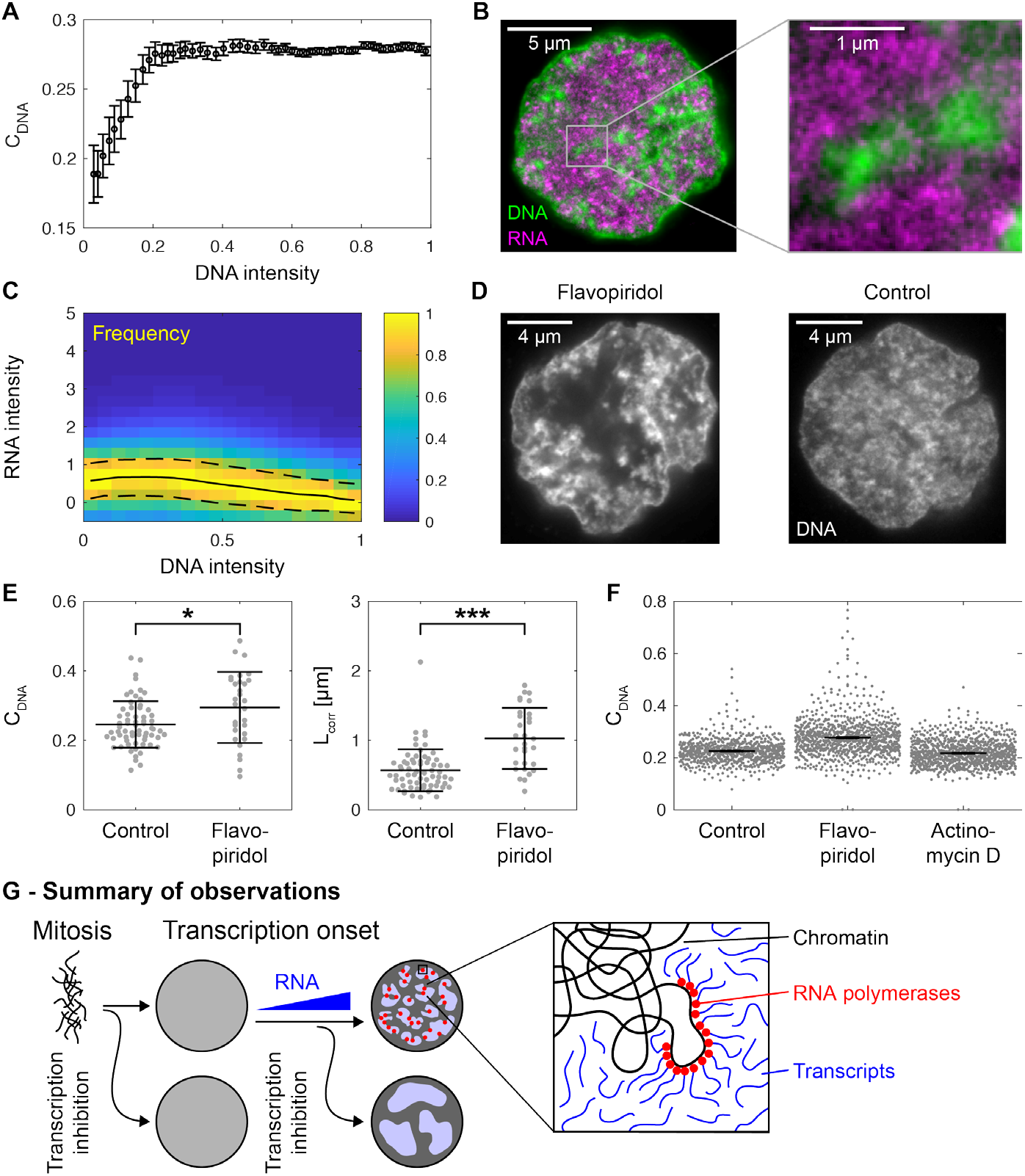
RNA accumulation establishes euchromatin domains, which are maintained in a finely dispersed pattern by transcriptional activity. **A)** Image contrast from nuclei of flavopiridol-inhibited cells, cells binned by nuclear RNA intensity, data recorded by spinning disk confocal microscopy (mean±s.e.m., analysis over 1582 cells in total). **B)** Representative STED micrograph showing spatial segregation of DNA and RNA in a nuclear mid-section from a flavopiridol-inhibited cell. **C)** RNA intensity distributions in flavopiridol-inhibited cells resolved by DNA intensity (solid line: median, dashed line: quartile range). **D)** Representative STED micrographs showing DNA intensity profiles in mid-sections of a control- and a flavopiridol-treated cell. **E)** Image contrast and correlation length after control and flavopiridol treatment (mean±std.dev., *p<0.05, ***p<0.001, permutation test, n=66, 30). **F)** Image contrast after treatment with different transcription inhibitors, recorded by spinning disk confocal microscopy (mean±s.e.m., n= 717, 886, 954, for cell selection see SI Figure 7). **G)** Sketch summing up experimental observations to this point.

Next, we studied the role of transcriptional activity in euchromatin organization. To exclude effects of variations in nuclear RNA level, we selected cells that retained significant amounts of nuclear RNA after inhibition with flavopiridol (SI Figure 4D). In the nuclei of these cells, RNA is localized to regions with low DNA intensity (Figure 2B,C), as was observed in nontreated cells. Hence, the segregation into chromatin domains and RNA-enriched regions appears to be unaffected by the suppression of transcriptional activity. The pattern formed by DNA domains, however, is markedly coarser in nuclei of inhibited cells when compared to nuclei of control cells (Figure 2D). DNA domains are more pronounced, as reflected by an increased DNA image contrast (Figure 2E, left panel), and larger, as reflected by an increased correlation length, a measure that quantifies the length scale of patterns in the DNA intensity profile (Figure 2E, right panel and SI Figure 6A, details see Supplementary Methods). The observed changes were not due to toxic side effects of flavopiridol treatment, as embryos treated with flavopiridol resumed normal development after washing out the drug (data not shown). Together, these observations suggest that transcriptional activity is required to maintain RNA and chromatin domains in a finely interspersed pattern.

We hypothesized that transcriptional activity stabilizes the finely interspersed pattern of RNA and chromatin domains by establishing physical contacts between these domains. Such contacts occur because transcribing RNA polymerases physically engage the DNA they are transcribing and simultaneously maintain a physical connection with the RNA transcript they are producing. If these contacts are sufficient to maintain a finely interspersed domain pattern, the active process of transcription would not be required. To test this prediction, we used the transcription inhibitor actinomycin D. In contrast to flavopiridol, which results in the loss of transcribing RNA polymerase from DNA (19), actinomycin D arrests RNA polymerases during transcription (20). The arrested polymerases are temporarily retained on DNA along with their associated transcripts (20). Indeed, actinomycin D treatment suppressed transcriptional activity in general (SI Figure 7), but a speckle pattern of polymerases that are retained on the DNA could still be observed (SI Figure 8). As predicted, no detectable coarsening of the DNA domains occurred in this case (Figure 2F). These results imply that the physical contact between DNA and RNA that is established by RNA polymerase, but not the process of transcription itself, is required to maintain the interspersed pattern of DNA and RNA domains.

Our experimental results to this point are summarized in Figure 2G. In brief, transcription onset after mitosis establishes a finely interspersed pattern of mutually exclusive chromatin domains and RNA-enriched regions. This interspersed pattern is maintained by DNA-RNA contacts established via RNA polymerases. When transcription is inhibited in a manner that removes these DNA-RNA contacts, chromatin domains and RNA-enriched regions are no longer finely interspersed. Instead, large scale demixing is observed. In nuclei not containing a significant amount of RNA, chromatin domains do not form, irrespective of the application of a transcription inhibitor.

To explain our experimental observations by a general principle, we devised a physical model of euchromatin organization by transcription. This model follows two main components: RNA-binding proteins (RBPs) and chromatin. We included RBPs in the model because RNA transcripts typically occur as RNA-RBP complexes (often referred to as ribonucleoproteins, RNPs) (21). In addition, RBPs are known to associate with RNA to form nuclear domains, such as splicing speckles and nucleoli (22). Moreover, membrane-less compartmentalization often arises from protein/RNA interactions (23). In our model, RBPs occur in two forms: unbound or bound to RNA in the form of RNA-RBP complexes. Unbound RBPs intermix with chromatin, whereas RNA-RBP complexes segregate from chromatin (Figure 3A, macromolecular mechanism 1). In agreement with this macromolecular mechanism, we found that the canonical splicing speckle protein SC35 mixed with DNA in nuclei with low amounts of RNA, but segregated from DNA in nuclei with high amounts of RNA (SI Figure 9). The second component of our model, chromatin, also occurs in two states: transcribed or nontranscribed. While chromatin generally segregates from RNA-RBP complexes, transcribed chromatin is retained in RNA-RBP complex-rich regions (Figure 3A, macromolecular mechanism 2). This is a consequence of the tethering of RNA-RBP complexes to chromatin during the transcription process. A detailed description of the model and parameter choice can be found in the Supplemental Text.

**Figure 3.**
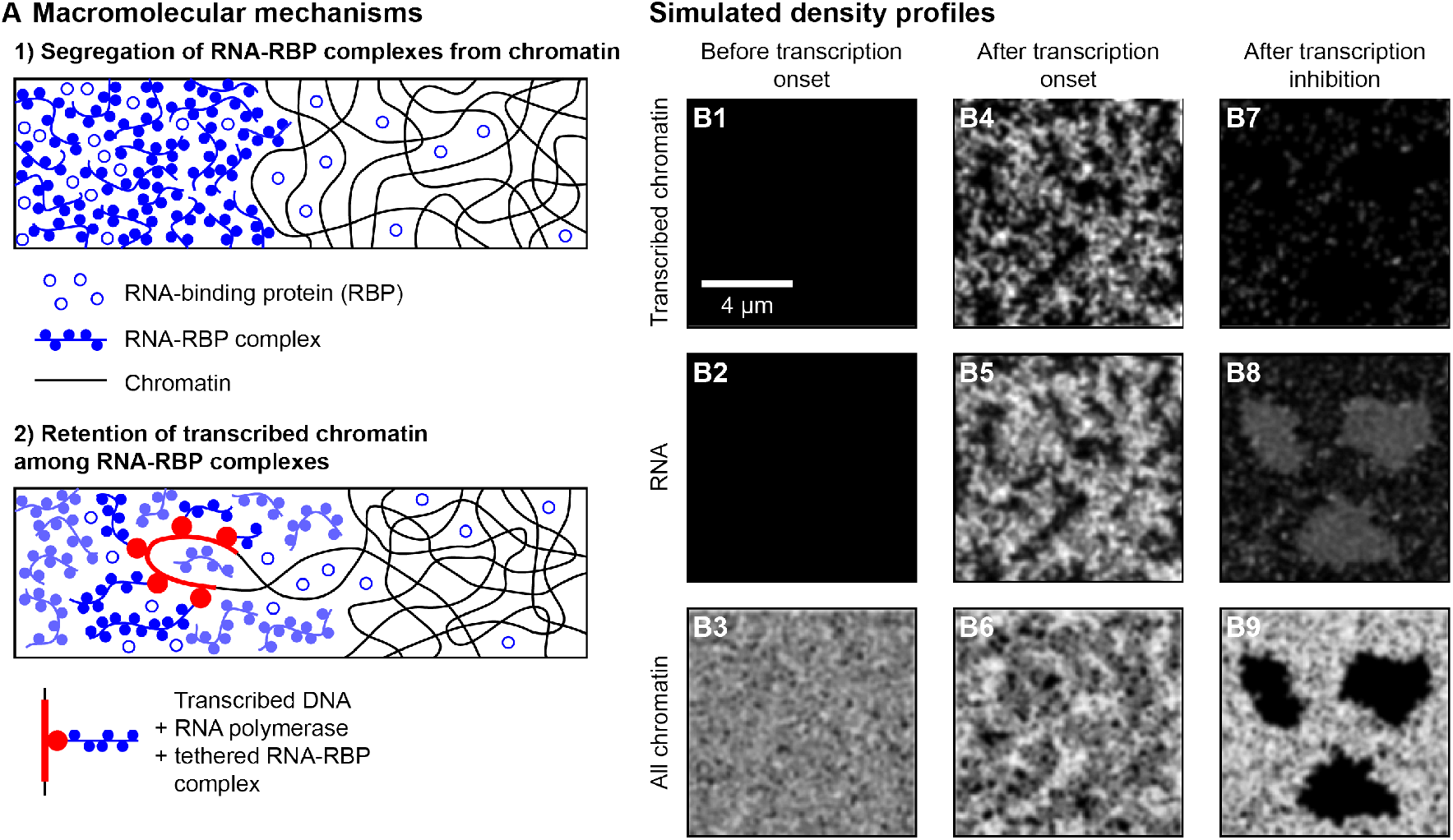
Physical model reproduces key features of euchromatin organization. **A)** Two macromolecular mechanisms were assumed in the construction of the model, for details see Supplementary Methods. **B)** Density profiles obtained from simulations of the physical model, approximating the conditions in a nucleus before transcription onset (B1-B3), after transcription onset (B4-B6), and following subsequent transcription inhibition (B7-B9). The densities of transcribed chromatin, RNA, and all chromatin are extracted from simulations in a way that they can be visually compared to transcriptional activity (Pol II Ser2Phos), RNA, and DNA in micrographs obtained in our experiments, respectively.

To test if our physical model can account for the DNA intensity profiles we observed in our experiments, we approximated three experimental conditions: before transcription onset, after transcription onset, and after transcription inhibition with flavopiridol (Supplementary Methods). Before transcription onset, neither transcribed chromatin nor RNA were present, and the chromatin concentration profile exhibited no domains (Figure 3B1-3). After transcription onset, both transcribed chromatin as well as RNA were present (Figure 3B4,5). Under these conditions, the chromatin concentration profiles exhibited domains which were interspersed with RNA accumulations and transcribed chromatin (Figure 3B6). Upon transcription inhibition, a significant amount of RNA was retained while most chromatin returned to the non-transcribed state (Figure 3B7,8). Here, the chromatin pattern was markedly coarsened when compared to the uninhibited situation (Figure 3B9). In all three cases, the simulated chromatin concentration profiles exhibited patterns similar to those observed in our experiments. Quantitative analysis of the chromatin density profiles also showed a good agreement between experiments and simulations (SI Figure 6B-D). Thus, a physical model can explain the key features of the euchromatin organization observed in our experiments.

Finally, we wanted to understand the dynamic process of microenvironment formation. Our work so far has revealed that microenvironments are central to euchromatin organization, but the sequence of events that underlies microenvironment formation and the resulting chromatin reorganization remains elusive. To investigate this sequence, we first simulated transcription onset at an isolated transcription site (SI Figure 10). We found that the onset of transcriptional activity is followed by the accumulation of RNA-RBP complexes around the transcription site. A DNA-depleted region is dynamically established as the accumulating RNA-RBP complexes displace non-transcribed chromatin, while transcribed chromatin is retained within the chromatin-depleted region. Next, to visualize the dynamic process by which transcription organizes euchromatin *in vivo*, we followed two prominent transcription sites that precede nucleus-wide transcription in practically all nuclei of late blastula zebrafish embryos (SI Figure 11A). These foci emanate from the repetitive microRNA miR-430 cluster (SI Figure 11B) (17,24), which is highly transcribed in early embryonic development (25). We used live cell-compatible antibody fragments that detect elongating RNA polymerase II (26,27) and cultured cells in refractive index-matched medium (28). Full transcriptional activity at the two prominent foci was established within a few minutes after mitosis (Figure 4A). At exactly the sites where, and the time when, we detected transcriptional activity, DNA was displaced (Figure 4A). Spatiotemporal analysis across multiple nuclei indicated that this sequence of events is highly reproducible (Figure 4B). Fixed cell microscopy confirmed that RNA accumulates with increasing transcriptional activity, and transcribed DNA is retained within the newly forming microenvironments (SI Figure 12). Hence, in model simulations as well as in live cells, RNA accumulation in the vicinity of transcription sites displaces nontranscribed DNA, while retaining transcribed DNA in the RNA-enriched microenvironment. In summary, our physical model explains both the nucleus-wide euchromatin organization as well as the formation of individual microenvironments observed in our experiments.

**Figure 4.**
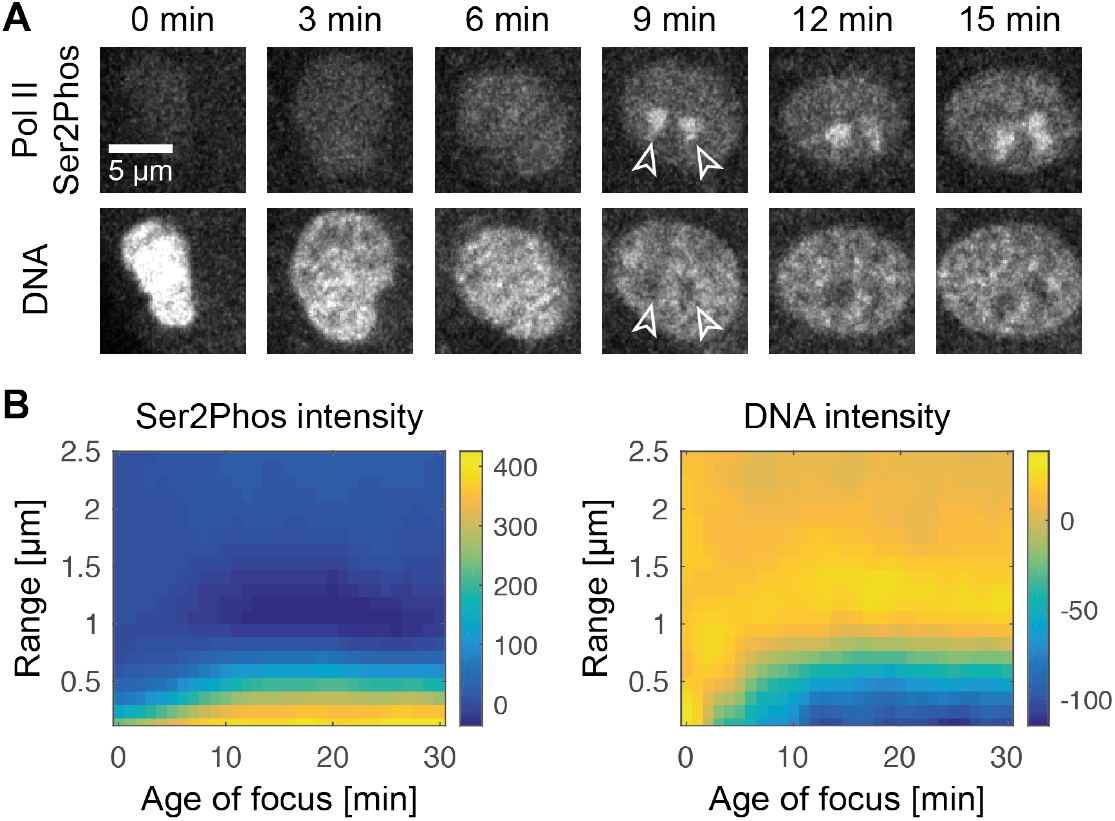
Transcription establishes microenvironments by dynamically replacing not-transrcibed chromatin. **A)** Representative time-lapse showing elongating RNA Polymerase II (Pol II Ser2Phos) and DNA in nuclear mid-sections following mitosis. Arrowheads indicate prominent transcription foci and zones of DNA depletion. **B)** Radial analysis, starting at the time when transcription foci were first detected. The range indicates radial distance from the centroid of a given Pol II Ser2Phos focus. Analysis averaged over 13 nuclei.

Our analysis of the 3D organization of euchromatin results in two major conclusions. First, we find that transcription establishes RNA-enriched microenvironments, which organize euchromatin. Transcription and RNA have been widely implicated in euchromatin organization before (29–31), but how they come together to organize euchromatin remained unclear. Our results show that RNA accumulates within localized microenvironments, and the exclusion of non-transcribed euchromatin from these microenvironments results in euchromatin domain formation. Conversely, active RNA polymerases tether RNA transcripts to the transcribed DNA, maintaining physical connections that stabilize the fine-grained pattern formed between RNA-enriched microenvironments and euchromatin domains. Together, this explains the widely conserved partitioning of transcribed and non-transcribed euchromatin into interchromatin space and chromatin domains, respectively (2), as well as the reduction of chromatin packing density in transcriptionally active euchromatin (32,33). In addition, our findings might explain how transcription can establish local microenvironments, where RNA polymerase and transcription factors accumulate and which have been proposed as spatial hubs in gene regulation (34–41). Finally, microenvironments provide a natural explanation for long range DNA-DNA contacts that frequently occur between highly transcribed elements (30,42–44). Together, this suggests that microenvironments play an important role in the spatial organization of transcriptional activity in the nucleus.

Second, returning to our search for a physical principle, we can categorize the 3D organization of euchromatin as an active microemulsion. Conventional microemulsions consist of two phases, often a hydrophobe and a hydrophile, and an amphiphile with affinity for both phases (45). The amphiphile, for example a detergent, stabilizes an interspersed pattern between the two phases. The segregation of RNA from non-transcribed chromatin, as seen in our experiments, suggests that RNA and non-transcribed chromatin correspond to the two phases of a microemulsion. Following this logic, the tethering of transcripts to chromatin via RNA polymerase II would result in the formation of an effective amphiphile with valencies for both phases. As expected for a microemulsion (45), the dissociation of these amphiphiles by RNA polymerase II inhibition results in coarsening of the pattern formed between the two phases. Differently from a conventional microemulsion, the amphiphile in our system synthesizes RNA transcripts, which convert freely diffusing RBPs into RNA-RBP complexes that segregate from chromatin. Hence, euchromatin organization can be described as an active microemulsion, which is stabilized by amphiphiles that also produce one of the phases. An increasing number of studies successfully applies physical principles to subcellular organization (46,47), indicating that often physical properties of the involved molecules, rather than their specific identities, are essential to establish spatial organization.

In the case of three-dimensional genome organization our work, together with previous experiments, indicates that heterochromatin is segregated from euchromatin by phase separation (14,15,48), while transcription internally organizes euchromatin similar to an active microemulsion.

## Acknowledgments

This work was supported by the Max Planck Society, a Human Frontier Science Program career development award (to N.L.V., CDA-00060/2012), an ELBE Postdoctoral Fellowship from the Center for Systems Biology Dresden (to L.H.), the Volkswagen foundation (to V.Z.), JSPS KAKENHI Grants JP15K07157 (to Y.S.) and JP25116005, JP26291071, and JP17H01417 (to H.K.). We thank Antonius van Boxtel for *in situ* probes, Jan Brugues, Anthony Hyman, Shai Joseph, Máté Pálfy, Iain Patten, Wolfram Poenisch, Jaques Prost, Thomas Quail, Rabea Seyboldt, Carine Stapel, and Christoph Weber for discussions, Stephan Grill, Carl Modes, Jochen Rink, and Christof Zechner for manuscript comments, and the following facilities and services: fish facility (MPI-CBG), light microscopy facility (MPI-CBG), computer department (MPI-CBG and MPI-PKS).

## Supplementary Materials

Materials & Methods

SI Figs. 1-12

## Supplementary Materials & Methods

### Embryo dissociation and cell culture

Wild type zebrafish (TLAB) were maintained and raised under standard conditions. Embryos were obtained by natural mating. Embryos were dechorionated within 20 minutes of fertilization and kept at 28.5°C. For dissociation into single cells, embryos in the late Oblong stage were immersed in 1 ml of deyolking buffer (10% v/v glycerol/H_2_O with 55 mM NaCl, 1.75 mMKCl, 1.25 mM NaHCO_3_) in low retention microcentrifuge tubes and vortexed at low speed until no intact embryo fragments could be observed. After centrifugation (1 min, 300 g), supernatant was aspirated and replaced with wash buffer (10% v/v glycerol/H_2_O with 110 mM NaCl, 3.5 mM KCl, 2.7 mM CaCl_2_, 10 mM Tris/Cl, pH 8.5), and tubes were vortexed at low speed to dissolve the cell pellet. After centrifugation (1 min, 300 g), supernatant was aspirated and replaced with 1 ml of PBS (all PBS in this study was Dulbecco’s formulation) with 0.8 mM CaCl_2_ added. Cells were cultured in this suspension for 30 min unless a different time is indicated. At the beginning of the time in suspension culture, tubes were briefly vortexed at low speed and then transferred into a rotator to prevent pellet formation.

### Transcription inhibition

#### α-manitin

α-manitin (A2263, Sigma) was dissolved and diluted to 0.2 mg/ml in H_2_O, and 1 nl of this solution was injected into embryos at the single cell stage to deliver 0.2 ng of α-manitin (Lee *et al.*, 2013). Control embryos were injected with 1 nl of H_2_O.

#### Flavopiridol

Flavopiridol (F3055, Sigma) was dissolved to 12.5 M (5 mg/ml) in DMSO, and diluted in PBS+0.8 mM CaCl_2_ to a final concentration of 1 μM for the application in suspension cell culture. Control cell cultures were kept in PBS + 0.8 mM CaCl_2_, with corresponding DMSO concentration. To test if the effect of flavopiridol was reversible, we assessed its effect on embryonic development. Embryos raised in 0.3X Danieu’s medium with 1 μM flavopiridol showed the typical developmental arrest before gastrulation. Normal development was resumed when the drug was washed out within an hour after arrest: embryos showed unperturbed muscle twitching, heartbeat, blood circulation, pigmentation, and swimming behavior during their further development.

#### Actinomycin D

Actinomycin D (A1410, Sigma) was dissolved to 1 mg/ml in DMSO, and diluted in PBS+0.8 mM CaCl_2_ to final concentrations of 5 μg/ml for the application in suspension cell culture. Control cell cultures were kept in PBS + 0.8 mM CaCl_2_, with corresponding DMSO concentration. Because it is known that actinomycin D is largely irreversible, we did not test reversibility.

### Fixed sample microscopy

#### Preparation of fixed cells for fluorescence staining

To compact the cultured cells into a pellet, suspension cultures were centrifuged during the last minute of cell culture (300 g). To fix the cells without perturbing the pellet, 8% formaldehyde in 1x PBS was added to the cell culture medium in a volume ratio of 1 in 4, to give an effective concentration of 2% formaldehyde. After 30 minutes of fixation at room temperature, tubes were centrifuged (1 min, 600 g), and supernatant aspirated. To increase the mechanical stability of cells, a secondary fixation step was carried out by applying 8% formaldehyde in PBS for 30 minutes at room temperature, followed by centrifugation (1 min, 800 g) and aspiration. To permeabilize the cell membrane and the nuclear envelope, 0.5% Triton X-100 in PBS was applied for 10 minutes at room temperature, followed by three washes with PBS with 0.1% Tween-20 (PBST).

#### Immunofluorescence labeling

Immunofluorescence labeling started with blocking samples in 4% (w/v) BSA in PBST for 30 min at room temperature. Primary antibodies were diluted in 2% (w/v) BSA in PBST and left to incubate at 4°C overnight. This was followed by three PBST washes at room temperature and subsequent application of fluorophore-conjugated secondary antibodies in the same way as the primary antibodies.

#### Total zygotic RNA labeling

Total zygotic RNA was labeled using the Click-iT RNA labeling kit (C10330, ThermoFisher). 1 nl of 50 μM 5-Ethynyl Uridine (EU, diluted from 100 μM stock in H_2_O) was injected into the cytoplasm of the first cell following fertilization, so that transcripts produced from the one-cell stage on incorporated EU. Click labeling of incorporated EU with an Alexa-594 azide was carried out following the manufacturer instructions, applying 100 μl click labeling mix per microcentrifuge tube. When combined with immunofluorescence staining, click labeling was carried out after permeabilization and before BSA blocking.

#### FISH labeling of primary miR-430 transcripts

miR-430 primary transcripts were labeled by RNA fluorescence *in situ* hybridization (RNA FISH) using a primary transcript probe kindly provided by Antonius van Boxtel (van Boxtel *et al.*, 2015). FISH probes were *in vitro* transcribed from linearized pGEMt_miR-430_ISH plasmid (Ndel restriction enzyme, New England BioLabs) using T7 polymerase (*in vitro* transcription mix: 2 μl transcription buffer (Roche), 2 μl DIG RNA labeling mix (Roche), 2 μl DTT stock (0.1 mM stock concentration), 1 μl RNAse inhibitor (Roche), 8 μl linearized DNA, 4 μl nuclease-free H_2_O, 1 μl T7 polymerase (produced in-house at Max Planck Institute of Molecular Cell Biology and Genetics), left to incubate at 37°C for 2 hours). *In vitro* transcription was followed by addition of 1 μl Turbo DNase (Ambion), incubation at 37°C for 1 hour, clean-up with QiaGen RNeasy MinElute Cleanup kit, and dilution in hybridization buffer (500 ml formamide; 65 ml 20X SSC, pH 5,0; 10 ml EDTA 0.5 M; 50 mg Torula yeast; 2 ml of 10% Tween-20 (v/v); 5 ml of 20% SDS (manufacturer stock concentration); 2 ml of 50mg/ml Heparin stock, filled up to 1 l, aliquoted to 50 ml, and stored at -20°C) to 50 mg/ml. The FISH procedure was started with one wash 50%/50% (v/v) Methanol/PBST, followed by two washes with 100% Methanol, then samples were placed at -20°C overnight. After returning samples to room temperature, two washes in 50%/50% Methanol/PBST and 2 washes in PBST followed. 70°C prewarmed hybridization buffer was added and samples were incubated for 1 hour at 70°C. Samples were then incubated in prewarmed hybridization buffer with 1:25 hybridization probe for 4 hours at 70°C, followed by three washes in hybridization buffer (70°C, 20 min each), one wash in 50%/50% (v/v) Methanol/PBST (70°C, 15 min exact), and three washes in PBST (room temperature, 10 min each), and 5% (v/v) blocking buffer (2% blocking reagent (Roche, 1 096 176) in 1X maleate buffer; maleate buffer: 150mM maleic acid, 100mM NaCl, pH 7.5, filter-sterilized, stored at room temperature) in PBST (room temperature, 20 min). Primary antibody incubation (mouse IgM monoclonal anti-Pol II Ser2Phos; Anti-Digoxigenin-POD, sheep Fab fragments) was in 2% BSA in PBST at 4°C for 48 hours, followed by three washes with PBST. FISH probes were revealed using the TSA Plus Cyanine 3 signal amplification kit (Perkin-Elmer), preparing 1 μl Cy3-Tyramide in 25 μl amplification buffer per sample, which was applied for 30 min at room temperature, followed by one wash in PBST. Incubation with secondary antibody (anti-mouse IgM-Alexa 488) was in 2% BSA in PBST at 4°C overnight, followed by three washes in PBST.

#### DNA labeling and mounting

DNA was labeled with DAPI or SiR-DNA (SC007, Spirochrome). DAPI was used for spinning disk confocal microscopy. DAPI was added directly into mounting media immediately before mounting at a concentration of 2 μg/ml. DAPI-stained samples were mounted in VectaShield H-1000, a non-setting liquid mounting medium. SiR-DNA was used for STED microscopy, RNA FISH labeled samples (FISH procedure induced high background on DAPI channel), and spinning disk confocal microscopy (equal or superior performance compared to DAPI). SiR-DNA staining produced no or very low signal in PBS, PBS+DABCO, or VectaShield H-1000, but signal was extremely bright when samples were mounted in glycerol-rich media. For this reason, SiR-DNA stained samples were mounted in glycerol. Because glycerol induced dissociation of several antibody combinations from the samples, immunofluorescence staining in these samples was followed by a post-fixation step of 30 min in PBS with 4% formaldehyde, 3 washes in PBST, and a careful but thorough replacement of PBST with ~20 μl of pure glycerol. We then diluted the SiR-DNA stock (1 mM in DMSO) in glycerol of which we spiked 1 μl into every sample immediately before mounting. The dilution of SiR-DNA in glycerol was adjusted so that upon addition to the 20 μl mounting medium the desired dilution was reached (1:60 in all cases except after α-manitin treatment, where 1:400 was used; all dilutions produced sufficient signal). Samples were mounted by spotting of mounting medium with resuspended cells onto regular microscope slides, applying #1.5 coverslips, and sealing with nail polish.

#### STED super-resolution microscopy of fixed cells

Measurements were performed on a commercial confocal STED microscope (Abberior Instruments, Göttingen, Germany) with pulsed laser excitation (490 nm, 560 nm, 640 nm, 40 MHz), beam scanning module (line frequency 3 kHz), a pulsed STED laser (775 nm, 40 MHz, spatial light modulator to produce the donut) and single photon counting APD detectors. Multicolor STED imaging with the single 775 nm STED laser was done by using chromatic separation of the fluorophores in combination with line-interleaved (time) excitation and detection. For the 560 nm and 640 nm channels, we used the dyes Alexa 594 and SiR, respectively. For the 490 nm channel, we used the long Stokes shift dye Abberior STAR 470 SXP, which emits in the 560 nm and 640 nm channel and can be effectively depleted by the 775 nm STED laser. To account for direct excitation of the SiR dye by the STED laser, we recorded the 640 nm channel additionally with only the STED laser activated. This channel was then subtracted from the SiR 640 nm channel.

#### Spinning disk confocal microscopy of fixed cells

Fixed cells were imaged using the Andor Revolution platform with Borealis extension, equipped with an Olympus silicone oil immersion objective (UPLSAPO 100XS, NA 1.35), recording with a single iXon Ultra 888 EMCCD camera. Acquisition settings were kept consistent across the different samples of a given experiment.

#### Light sheet imaging of whole fixed embryos

Fixed whole embryos were prepared, fluorescently stained, and imaged using a Zeiss Z1 light sheet microscope exactly as described by us in a previous publication (Joseph *et al.*, 2017). Pol II Ser2Phos was labeled by immunofluorescence, using mouse IgM anti-Pol II Ser2Phos primary antibody (1:500) and anti-mouse IgM secondary antibody (conjugated with Alexa 488, dilution 1:1000). DNA was stained by adding 1 μg/ml DAPI during secondary antibody incubation.

#### List of antibodies

##### Primary antibodies

Mouse IgM anti-Pol II CTD Ser2Phos (H5), monoclonal, ab24758 abcam

Dilution: 1:500 for light sheet microscopy

Rabbit IgG anti-Pol II CTD Ser2Phos, monoclonal, ab193468 abcam

Dilutions: 1:200 for STED microscopy, 1:1000 for confocal microscopy

Rat IgG anti-H3 Ser28Phos, monoclonal, ab10543, abcam

Dilution: 1:1000 for confocal microscopy

Sheep IgG Anti-Digoxigenin Fab fragments, conjugated with horseradish peroxidase, 1207733 Roche; Dilution: 1:500 for fluorescence in situ hybridization

Mouse IgG aSC35, monoclonal, 556363 BD Biosciences

Dilution 1:100 for confocal microscopy

##### Secondary antibodies

Goat anti-mouse IgM, conjugated with Alexa 488, A21042 Thermo Fisher

Dilution: 1:1000 for light sheet microscopy

Goat anti-rabbit IgG, conjugated with STAR 470 SXP, 2-0012-008-9, Abberior

Dilution: 1:200 for STED microscopy

Donkey anti-rabbit IgG, conjugated with Alexa 488, A21206 Thermo Fisher

Dilution 1:1000 for confocal microscopy

Donkey anti-rat IgG, conjugated with Alexa 488, A21208 Thermo Fisher

Dilution 1:1000 for confocal microscopy

Goat anti-rat IgG, conjugated with Alexa 647, A21247 Thermo Fisher

Dilution 1:1000 for confocal microscopy

## Live cell microscopy

### Preparation of antibody fragments for use in live cell microscopy

The fluorescently labeled antibody fragments (Fabs) specific to Pol II Ser5Phos and Pol II Ser2Phos were prepared as described previously (Stasevich, Hayashi-Takanaka, *et al.*, 2014; Kimura and Yamagata, 2015). Briefly, monoclonal antibodies specific to Pol II Ser5 and Ser2 phosphorylations were digested with Ficin (ThermoFisher Scientific) and Fabs were purified through protein A Sepharose columns (GE Healthcare) to remove Fc and undigested IgG. After passing through desalting columns (PD MiniTrap G25; GE Healthcare) to substitute the buffer with PBS, Fabs were concentrated up to >1 mg/ml using 10 k cut off filters (Amicon Ultra-0.5 10 k; Merck), Fabs were conjugated with Alexa Fluor 488 (Sulfodichlorophenol Ester; ThermoFisher Scientific) or Cy3 (N-hydroxysuccinimide ester monoreactive dye; GE Healthcare) to yield ~1:1 dye:protein ratio. After the buffer substitution with PBS, the concentration was adjusted to ~1 mg/ml.

### Preparation of live cells for fluorescence microscopy

Directly following fertilization, zebrafish embryos were pronase-dechorionated and 1 nl of a mix made up of 0.3 μl Alexa 488-conjugated Pol II Ser5Phos Fab, 1.7 μl Cy3-conjugated Pol II Ser2Phos Fab, 0.2 μl 1 mM SiR-DNA, and 0.1 μl 10x Phenol Red was injected into the cytoplasm at the single cell stage. Embryos were grown at 28.5°C and dissociated into single cells at High stage. These cells were mounted in refractive index matched medium exactly as previously described (Boothe *et al.*,2017). During the time required to mount the cells and start microscopy, cells had undergone one to two divisions. In intact embryos, cells also undergo one or two cell divisions during the developmental progression from High to Oblong or Sphere stage. Thus, we acquired live microscopy images from cultured cells that should most closely correspond to cells at the Oblong or Sphere stage in the intact embryo.

### Confocal microscopy of live cells

Live cell cultures were imaged using the Andor Revolution platform with Borealis extension, equipped with an Olympus silicone oil immersion objective (UPLSAPO 100XS, NA 1.35), recording with a single iXon Ultra 888 EMCCD camera. Image data were acquired for up to 4 cell clones in parallel. A full three-color z-stack could be obtained every minute for all cell clones. Time-lapses were recorded over periods of up to 90 minutes, during which cells continuously displayed cell divisions, suggesting no obvious phototoxicity.

## Image preparation and analysis

### Software used for image preparation and analysis

Microscopy image preparation was done using FIJI (Schindelin *et al.*, 2012) and MatLab, the latter relying on the Open Microscopy Environment plugin for image import (Goldberg *et al.*, 2005). Further data processing was carried out in MatLab. The resulting figures were prepared for publication using MatLab and Adobe Illustrator.

### Segmentation of nuclei

The nuclei in STED images are segmented by applying Otsu’s method for adaptive thresholding to the DNA channel. In some cases, the resulting segmentation mask contains holes, which are removed by a filling step. Distortion and artifacts from out-of-focus light are seen at the boundaries of nuclei. To remove these imaging imperfections from further structural analysis, the segmentation masks are eroded before further analysis.

Spinning disk confocal microscopy data contain several nuclei and consist of a stack of multiple images in the z direction. An initial segmentation step based on a fixed, manually chosen threshold is applied to the DNA channel to obtain substacks containing individual nuclei. To extract a single image close to the middle of the nucleus in a given stack, the z section with the highest intensity contrast in the DNA channel is selected for further analysis. In this image, the nucleus is segmented using the same approach as described for STED images above. Images from STED and spinning disk confocal microscopy can be analyzed in the same manner from here on.

### Calculation of nuclear intensities

The mean nuclear intensity of a given color channel is extracted using the nuclear segmentation masks obtained from the DNA channel. These mean nuclear intensities contain contributions from actual nuclear signal and also image background intensity. To remove image background intensity, the fluorescence in the cytoplasm is determined and subtracted from the total nuclear intensity. The cytoplasmic intensity is determined using a segmentation shell that is created by an outward dilation of the nuclear segmentation mask (Stasevich, Sato, *et al.*, 2014; Joseph *et al.*, 2017).

### Calculation of image contrast

The DNA image contrast (*C_DNA_*) is calculated as the root mean square contrast of the individual pixels’ intensities (*I_n_*) and normalized by the mean intensity, *Ī* = 〈*I_n_*〉,

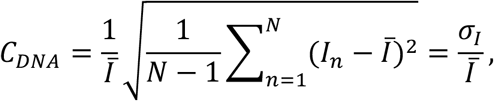

where *σ_I_* is the standard deviation. This is equivalent to the coefficient of variation of *I_n_*.

The *C_DNA_* of samples prepared, stained, and imaged under comparable conditions and identical settings can be quantitatively compared. To compare between images obtained under different conditions, background intensity correction is required. This is also required when microscopy images and simulated chromatin concentration profiles. An appropriate background correction can be calculated assuming an offset to the individual intensity values,

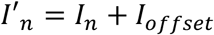

This leads to a changed image contrast value,

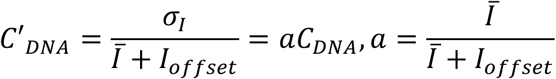

Thus, assuming of a constant *I_offset_*, *C_DNA_* values obtained under different conditions can be normalized by a reference condition, and relative changes in *C_DNA_* can be compared. This approach is used to compare between STED microscopy images and chromatin concentration profiles obtained from simulations. Specifically, the condition “after transcription onset” is used to establish the value of *a*, and the *C_DNA_* values obtained from simulations are divided by *a* before comparison.

### Calculation of correlation length

The correlation length of the DNA intensity distribution (*L_corr_*) is determined in two main steps. First, the radial correlation function, *g(r)*, is extracted. We use a definition of the radial correlation function that takes into consideration the segmentation mask covering the inside of the cell nucleus. Considering a DNA pixel intensity image *I_i,j_*, with the two-dimensional position of the pixel indicated by *i* and *j*, and an associated segmentation mask *σ_i,j_*, ∈ {0,1}, the radial correlation function at a distance *r* is

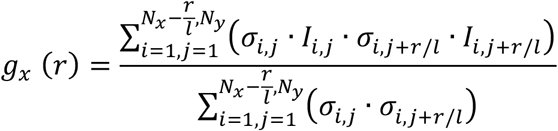

in the case of shifting in the x direction. Note that, due to the pixel resolution *l*, *g_x_* (*r*) is only evaluated at discrete intervals *r* = 0, *1*, 2*1*, &, *N_x_l*. The equivalent calculation is carried out for shifts in y direction to obtain *g_y_*(*r*). The combined radial correlation function then is *g*(*r*) = (*g_x_* + *g_y_*)/2. Before the calculation of *g(r)*, the intensities of all color channels are normalized by the respective color channels’ mean intensity in the segmented nucleus, followed by subtraction of the mean intensity in the segmented nucleus.

Second, to obtain *L_corr_*, an exponential decay function is fitted to *g(r)*. To this end, the function

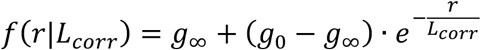

is adjusted to *g(r)* by optimization of the value of *L_corr_*. Here, *g*_0_ = *g*(*r* = 0) and *g*_∞_, representing the plateau level of the decaying correlation function, was approximated by the mean value of *g(r)* in the interval of *r* from 4.5 to 6.0 μm.

A common approach to structural characterization, Fourier analysis, cannot be used. Given that the structural analysis has to be contained to the inside of cell nuclei, domains with irregular boundaries need to be analyzed. It is not clear how Fourier analysis can be applied to such irregular domains in a straight-forward manner.

### Intensity distributions of one color channel with respect to another color channel

To determine the relationship between the intensity profiles of different color channels, we analyze the distribution of fluorescence intensities of a given channel (A) with respect to intensities in another color channel (B). To this end, all pixels of an image are binned based on the intensities of channel B. Then, the mean intensity on the channel A of all pixels within a given bin is calculated. This analysis reveals the intensity distribution of color channel A with respect to intensities in the color channel B.

The same principle can be applied to resolve a color channel A by the intensities of two other color channels, B and C. Instead of binning pixels only with respect to a single color channel (one-dimensional binning), the pixels are now binned with respect to two color channels (two-dimensional binning).

### Analysis of live cell images

At every time point, nuclei are segmented based on Pol II Ser5Phos Fab signal. Specifically, we first use the fact that signal of Pol II Ser5Phos Fab occurred in nuclei but also throughout the cytoplasm to segment cells from background using an Otsu threshold. Second, we use the higher signal intensity within nuclei to segment nuclei from cytoplasm, by applying an Otsu threshold within the segmented cells. When the Otsu metric is below 0.65, nuclei are segmented. Otherwise, it is assumed that the Fab pool was released to the cytoplasm due to nuclear envelope breakdown during mitosis, and no nuclei are segmented. For all pixels within segmented nuclei, their 3D distance to the nearest non-segmented pixel is calculated. To segregate nuclei that are too close to be directly segmented, a water-shed segmentation is initiated from the maxima of this distance map. The segmented nuclei are first automatically tracked through time by their centroid distance. Where tracks have gaps, or are not correctly connected, tracks are then manually corrected.

To analyze spatial organization around the two prominent transcription sites, we carried out a radial intensity analysis that is centered on these. Before any analysis, all fluorescence images are locally corrected for background intensity: each xy image is copied, filtered with a Gaussian kernel (kernel width of σ=2.38 μm), and subtracted from the unprocessed image. Transcription sites are segmented with an Otsu threshold applied to the Pol II Ser2Phos channel, and the two largest objects are retained, assuming that they are the two prominent transcription sites. For both these objects, the centroid is determined, and the xy-section containing the centroid is extracted for radial analysis. Within these xy-sections, the pixel containing the centroid is marked as the starting point of the analysis. With respect to the radial range of the analysis, this pixel is located at a range of 0, referring to the center of the transcription site. The first radial outward step now marks all 8 neighbors of this initial pixel, and refers to a radial range of 1 pixel. A radial range of 2 pixels is reached by marking the next line of outward-lying neighbors, and so forth for all further ranges. At all ranges, the mean intensities of Fab Pol II Ser5Phos, Fab Pol II Ser5Phos, and SiR-DNA signal within the pixels belonging to this radial range is calculated. This procedure produces an intensity curve for all color channels at different radial ranges with respect to the centroid of a given transcription site. To average over the transcription sites of several nuclei, the tracked nuclei were temporally aligned by the first time at which two transcription foci could be detected in a given nucleus. Two-dimensional images of intensity resolved by radial range and time were then created for each tracked nucleus. These were averaged over all tracked nuclei to create final plots.

## Physical model

### Model outline

We simulate the spatial organization and chemical conversions of macromolecular components (chromatin and RNA-binding proteins, RBPs, described in the main article) using a coarse-grained model, in which the space of interest is divided into discrete compartments. Each compartment is occupied by a single species, which represents the predominant component in this compartment. Coarse-graining makes simulations of large spaces computationally tractable, which would be too computationally intensive when simulated at the molecular level. To achieve acceptable computational performance in our case, we implemented our model as a two-dimensional square lattice. The simulation model used to implement the spatial organization is adapted from an approach originally used for microemulsions (Larson, Scriven and Davis, 1985). In brief, our approach allows individual compartments to undergo chemical conversions and allows neighboring compartments to stochastically swap contents. The likelihood of a chemical conversion is based on the rate of the respective reaction rate. The likelihood of a given swap is determined by the required free energy change. A free energy cost is associated with placing RNA-RBP complexes next to chromatin, representing the segregation of RNA-RBP complexes from chromatin. Note that, because RNA-RBP complexes are tethered to transcribed chromatin, this free energy cost also applies for the placement of transcribed chromatin next to chromatin in general. To account for the integrity of the linear DNA polymer, swap operations that would break chromatin into disconnected domains are not allowed.

### Detailed model implementation

We now describe our physical model in more detail (see Figure “Implementation of the physical model” below). Our model follows conversions of species as described by a chemical reaction network, as well as the spatial configuration in a twodimensional, square lattice with *N × N* sites (Model Figure 1). The chemical reactions are simulated using an iteration with a constant time step, 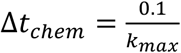, where *k_max_* is the maximal reaction rate that can occur in the reaction network. In every iteration step, the species present at a given lattice site can only undergo one reaction. Therefore, for every iteration step, for each site we compare a uniformly distributed random number *r*, with 0 ≤ *r* ≤ 1, to the reaction probability *P_reaction_* = Δ*t_chem_k_reaction_*, where *k_reaction_* is the rate of a given conversion reaction. When *r* ≤ *P_reaction_*, the reaction is executed.

**Model Figure 1:**
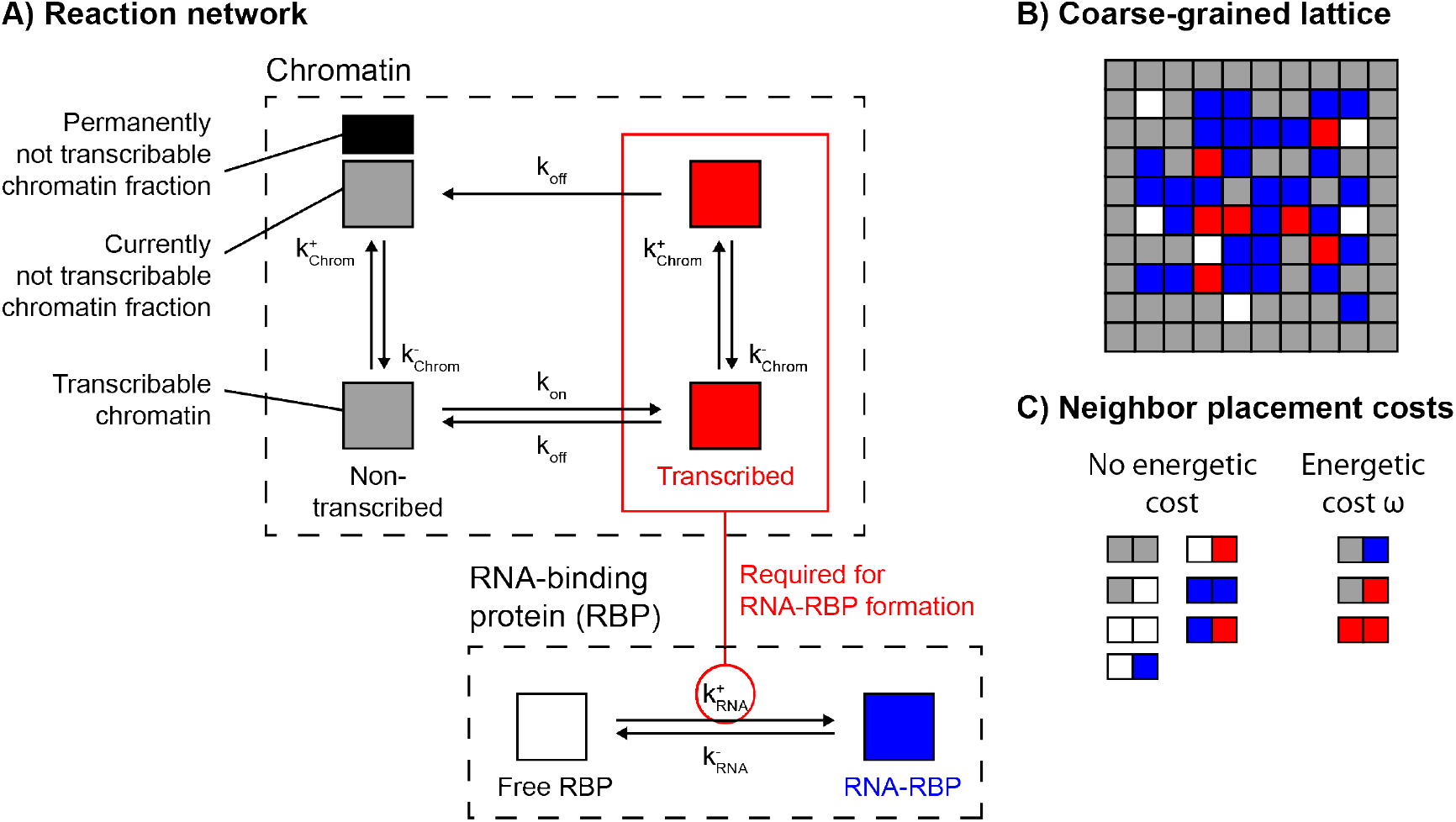
*Implementation of the physical model*. **A)** Reaction network describing the chemical conversions of individual lattice sites, rates indicated next to transitions. **B)** Representation of a square lattice used to follow the spatial configurations, here shown for *N* = 8. **C)** Overview of neighbor placements without or with an associated energetic cost (ω = 0.15 *k_B_T*).

Species located at different lattice sites change their location stochastically with direct neighbors. Swap operations are also simulated by iteration with a constant time step, Δ*t_lattice_*/*N*^2^, with Δ*t_lattice_* ≪ Δ*t_chem_*. The iteration for swap operations is interleaved with the iterations for chemical conversions. For every step in the iteration for the swap operations, a lattice site is randomly chosen, followed by random choice of a swap partner from the set of eight direct neighbors. For these two partners, the potential energy stored in the lattice configuration before and after the swap is calculated, Δ*E_swap_* = *E_post_* − *E_pre_*. *E_pre_* and *E_post_* are the free energies contained in the local neighbor pairings before and after the proposed swap operation, respectively. The probability of the swap to actually occur is

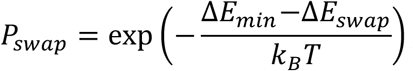

where Δ*E_min_* < 0 is the maximally possible free energy release in a single swap. The normalization by Δ*E_max_* ensures that 0 < *P_swap_* ≤ 1, and thereby the linearity of time in lattice reconfigurations. The actual free energies of local lattice configurations are calculated from neighbor configurations: every neighboring pair consisting of chromatin/RNA-RBP complex or chromatin/transcribed chromatin imposes an energy cost *w* > 0. Considering all possible neighbor configurations, the maximal free energy release from a swap then is Δ*E_min_* = −10*w*.

To ensure the polymeric integrity of chromatin, a connected components check is executed before every swap operation that involves a chromatin particle. The swap operation is aborted if a breaking of the overall chromatin into an increased number of connected components is detected. The lattice is padded around its margin by stationary chromatin, which allows proper energy calculations at the margins and represents chromatin anchored at the nuclear envelope. Also, swap operations that remove chromatin from the direct neighborhood of the padding chromatin layer were aborted, mimicking anchoring via envelope-attached chromatin.

In our model, the conversion of chromatin from the not transcribed to the transcribed state occurs in two steps. First, the chromatin is divided into subdomains, which represent contiguous gene bodies. These domains can switch between a not transcribable and a transcribable state. Second, when a given subdomain is in the transcribable state, the individual chromatin sites that are part of the subdomain can transition into the transcribed state (with rate 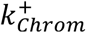). Transcribed chromatin sites can always transition back into the not transcribed state (with rate 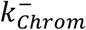). Subdomains are assigned when the simulation is initialized, by subdividing the chromatin into 1-by-1 μm^2^ subdomains. At the beginning of the simulation all subdomains are in the not transcribable state. The switching of subdomains into the transcribable state is implemented differently, dependent on the type of scenario that is simulated. In simulations at the whole nucleus level, initially the subdomains are kept off for a number of simulation steps corresponding to 5 minutes real time. Then, for a fraction of 60% of all domains, a non-zero rate of switching into the transcribable state is assigned (*k_on_*). At all times, the rate for a given subdomain to revert to the not transcribable state is kept at the same non-zero value (*k_off_*). To approximate transcription inhibition, all subdomains are again assigned a rate of zero to transition into the transcribable state. In simulations of transcription onset at an isolated transcription site, all subdomains but one are kept in the not transcribable state for the entire simulation. A single subdomain in the center of the lattice is assigned as transcribable at the beginning of the simulation. In all scenarios, all chromatin subdomains are also monitored as connected components throughout the simulation to maintain their polymeric integrity.

Compartments of unbound RBP are converted into RNA-RBP complexes only when one or more of the eight direct neighbor compartments contained transcribed chromatin, with a rate 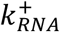. This dependence on transcribed chromatin is intended to represent the RBP binding of RNA transcripts produced locally as a result of transcription. RNA-RBP complexes are then either exported from the nucleus and replaced by unbound RBPs or are degraded over time inside the nucleus. Both processes are approximated by a constant decay rate 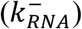 from RNA-RBP complexes to unbound RBP. To clarify, RNA is not explicitly treated in our model, but only indirectly monitored as a component of transcribed chromatin and RNA-RBP complexes. The total number of chromatin-containing compartments (transcribed as well as not transcribed) and the total number of RBP-containing compartments (unbound RBP as well as RNA-RBP complexes), however, are both conserved.

This model was implemented as C++ code (available for download at https://cloud.mpi-cbg.de/index.php/s/Mrt6nwt83jEEPvZ), which was compiled and executed on the computational cluster of the Max Planck Institute for the Physics of Complex Systems. Concentration profiles of total chromatin and transcribed chromatin were created directly from the respective simulation results. In our microscopy images, RNA signal resulted from a population of RNA not associated with ongoing transcription as well as from high intensity RNA foci that are closely associated with sites of ongoing transcription. We therefore calculated RNA concentration profiles by adding the RNA-RBP complex profile and the transcribed chromatin profile. Their relative contributions were weighted with a factor of 0.2 for the RNA-RBP complex profile and a factor of 0.8 for the transcribed chromatin. To convert the coarse-grained, “all-or-nothing” lattice simulation results into graded concentration profiles, all channels were blurred with a Gaussian filter (kernel width of σ=100 nm). To allow a comparison to the quantitative analyses of DNA organization in our experiments, we applied the same analysis procedures used for microscopy images also to the chromatin concentration profiles obtained from the above simulations. Model parameters are assigned from literature or chosen based on our experimental data (see Table “Model parameters” and Model Figure 2).

**Table:** Model parameters. Lowered relative to the value of *w*=0.5 k_B_T from (Larson, Scriven and Davis, 1985) to allow for a sufficiently rapid spatial reorganization in our simulations. ^2^Adjusted to achieve sufficient compaction after simulated inhibition compared to our experimental data. ^3^Estimated from our live imaging data, using the times from Pol II recruitment (Pol II Ser5Phos) to transcriptional activity (Pol II Ser2Phos). ^4^Rate of Pol II escape from promoter-paused state (Stasevich, Hayashi-Takanaka, *et al.*, 2014). ^5^Average time of 10 minutes for transcript completion estimated based on a typical length of genes transcribed at late blastula stage of 10 kb (Heyn *et al.*, 2014) and a typical Pol II transcription rate of 1 kb/min (Jonkers, Kwak and Lis, 2014; Stasevich, Hayashi-Takanaka, *et al.*, 2014). ^6^Within the range of typical nuclear retention times of completed transcripts (Bahar Halpern *et al.*, 2015; Battich, Stoeger and Pelkmans, 2015).

| Parameter | Property | Value |
| --- | --- | --- |
| $w$ | Reciprocal temperature | $0.15 \text{ k}_B\text{T}^1$ |
| $\Delta t_{\text{lattice}}$ | Swapping time step | $5 \times 10^{-5} \text{ min}^2$ |
| $k_{\text{on}}$ | Domain on rate | $0.075 \text{ 1/min}^3$ |
| $k_{\text{off}}$ | Domain off rate | $0.15 \text{ 1/min}^2$ |
| $k_{\text{Chrom}}^+$ | Transcription onset | $0.4 \text{ 1/min}^4$ |
| $k_{\text{Chrom}}^-$ | Transcription termination | $0.1 \text{ 1/min}^5$ |
| $k_{\text{RNA}}^+$ | RNA production rate | $0.5 \text{ 1/min}^2$ |
| $k_{\text{RNA}}^-$ | RNA decay rate | $0.05 \text{ 1/min}^6$ |

**Model Figure 2:**
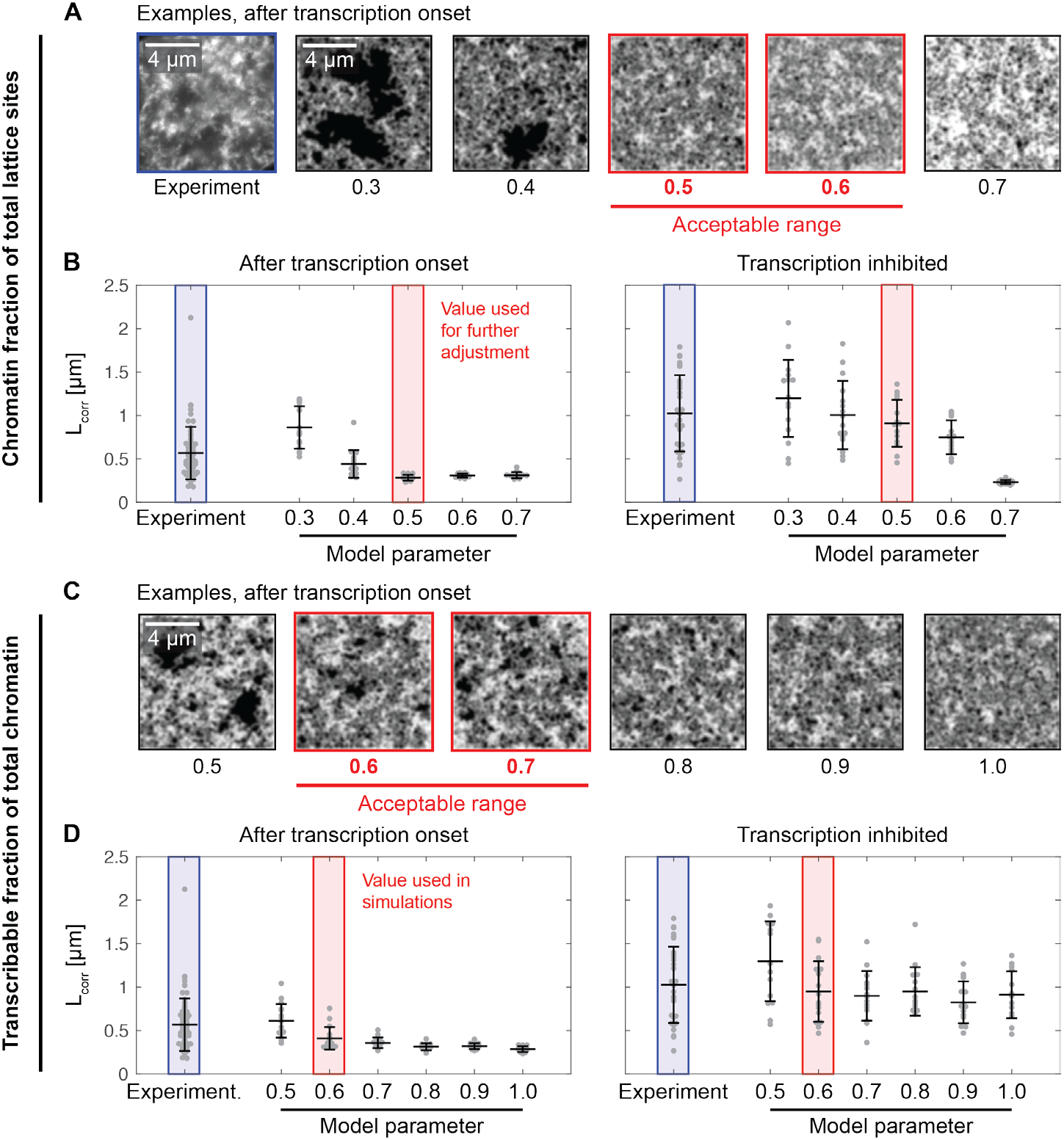
*Choice of total chromatin fraction and transcribable chromatin fraction in the physical model*. The total chromatin fraction and the fraction of chromatin that is transcribable in our coarse-grained model are parameters that cannot be directly inferred from literature or databases. Here, we execute simulations with different values of these model parameters and compare the resulting chromatin concentration profiles to our microscopy results. **A)** We execute simulations with different total chromatin fraction, while keeping the transcribable fraction at 1.0. A visual assessment of concentration profiles after transcription onset indicates that fractions of 0.5 and 0.6 are in acceptable. Below 0.5, large zones without any chromatin occur, which do not reflect the microscopy image. Above 0.6, the chromatin is too dense to permit formation of low chromatin concentration regions, which are observed in microscopy images. **B)** An assessment of the correlation lengths (*L_corr_*) of the chromatin concentration profiles from model simulations indicates that 0.5 as well as 0.6 would be acceptable chromatin fractions. We choose 0.5 as a value to use in our simulations. Values from 16 individual simulations are shown with mean±SD. **C)** To choose the transcribable fraction of total chromatin, we keep the total chromatin fraction of 0.5, and produce chromatin concentration profiles for different values of the transcribable fraction. Visual assessment indicates that transcribable fractions of 0.5 and 0.6 are acceptable. Below 0.5, large regions without chromatin occur, which do not agree with microscopy images. Above 0.7, the chromatin concentration profile looks too smooth compared to microscopy images. **D)** Lowering the transcribable fraction towards 0.5 brings *L_corr_* after transcription onset closer to experimental values, so that we choose the lowest acceptable transcribable fraction, which is 0.6. Values from 16 individual simulations are shown with mean±SD.

### Approximation of experimental conditions

To test if the physical model can account for the DNA intensity profiles observed in our experiments, we approximated experimental conditions and compared the resulting concentration profiles to our microscopy images (simulation results and comparison described in main article). First, to approximate the conditions of a cell before transcription onset, we set the rate at which chromatin transitioned into the transcribed state to zero and executed a number of simulations steps sufficient to equilibrate the system. Then, to approximate the conditions of a cell after transcription onset, we changed the rate at which chromatin transitioned into the transcribed state to a non-zero value. We continued the simulations until the concentration of RNA and transcribed chromatin reached a plateau. Lastly, to approximate transcription inhibition, we returned the rate at which chromatin transitions into the transcribed state to zero. We continued the simulations for a number of steps corresponding to 30 minutes, which was the duration of transcription inhibition in the experiment.

## Supplementary Figures

**SI Figure 1.**
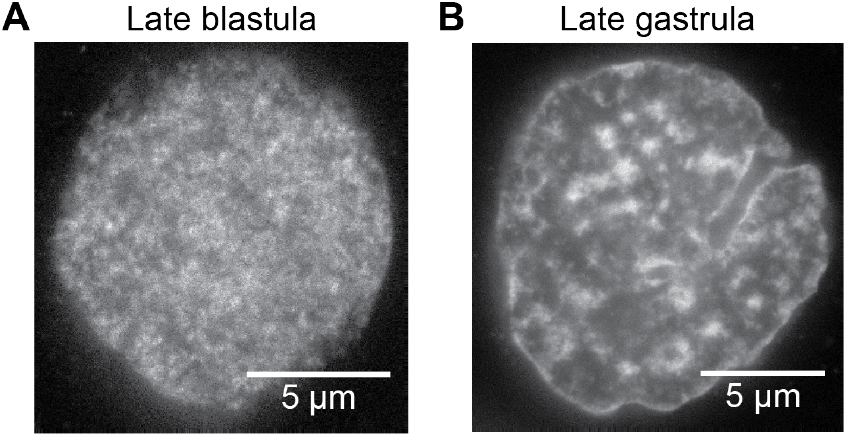
Heterochromatin domains are not observed in nuclei of late blastula zebrafish embryos. STED super-resolution micrographs from nuclear mid-sections in zebrafish late blastula embryos did not exhibit the highly compacted heterochromatin domains (A, sphere stage of development) that were present in late gastrula embryos (B, 80% epiboly stage).

**SI Figure 2.**
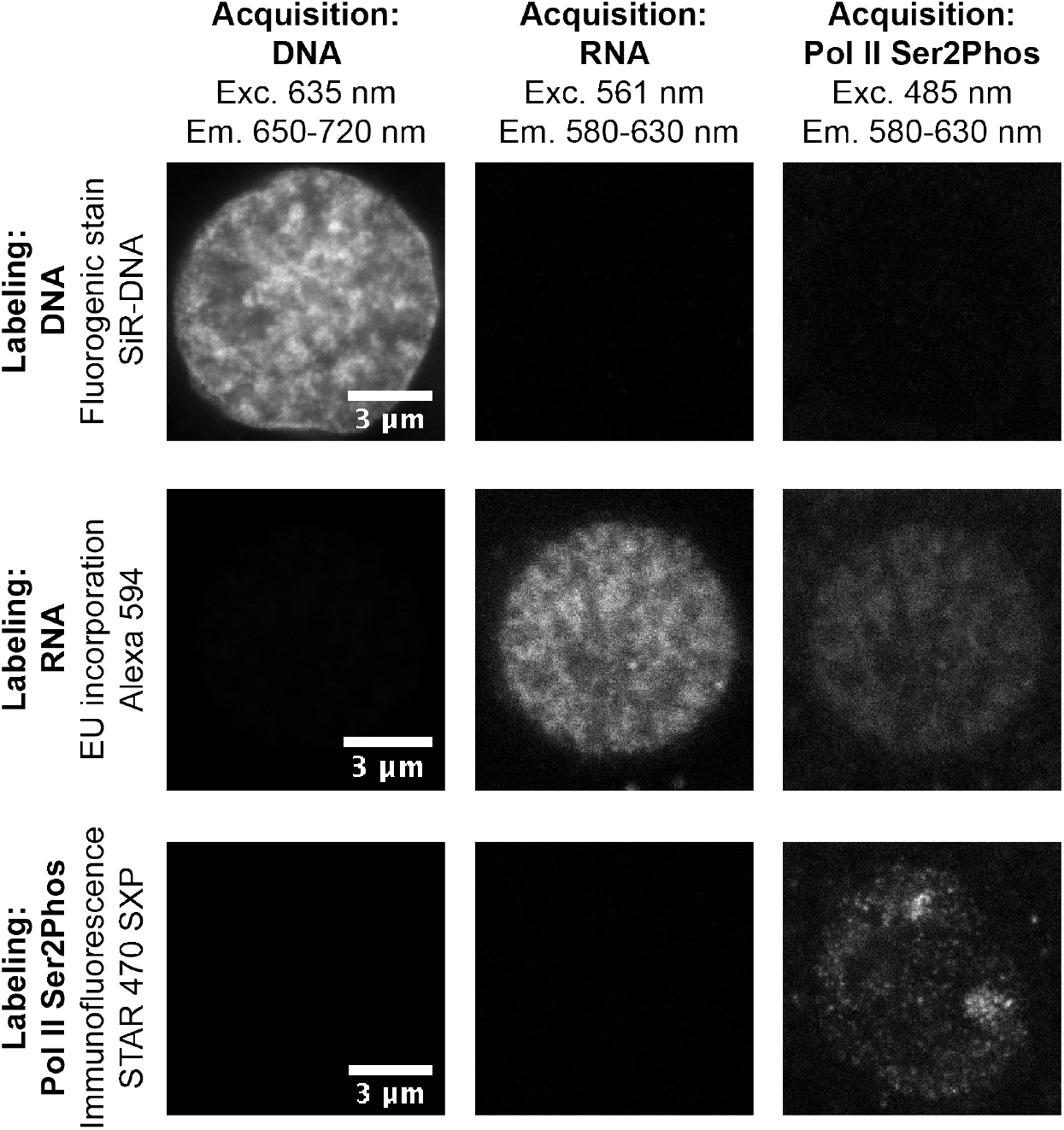
Three-color STED super-resolution microscopy of DNA, RNA, and transcriptional activity. Three STED super-resolution micrographs from nuclear midsections, stained for either DNA, RNA, or transcriptional activity (detected by RNA polymerase II C terminal domain Serine 5 phosphorylation, Pol II Ser2Phos). Each label can be detected in one color channel with negligible crosstalk to the other channels. For each label, we acquired three micrographs, and here show one representative micrograph per label.

**SI Figure 3.**
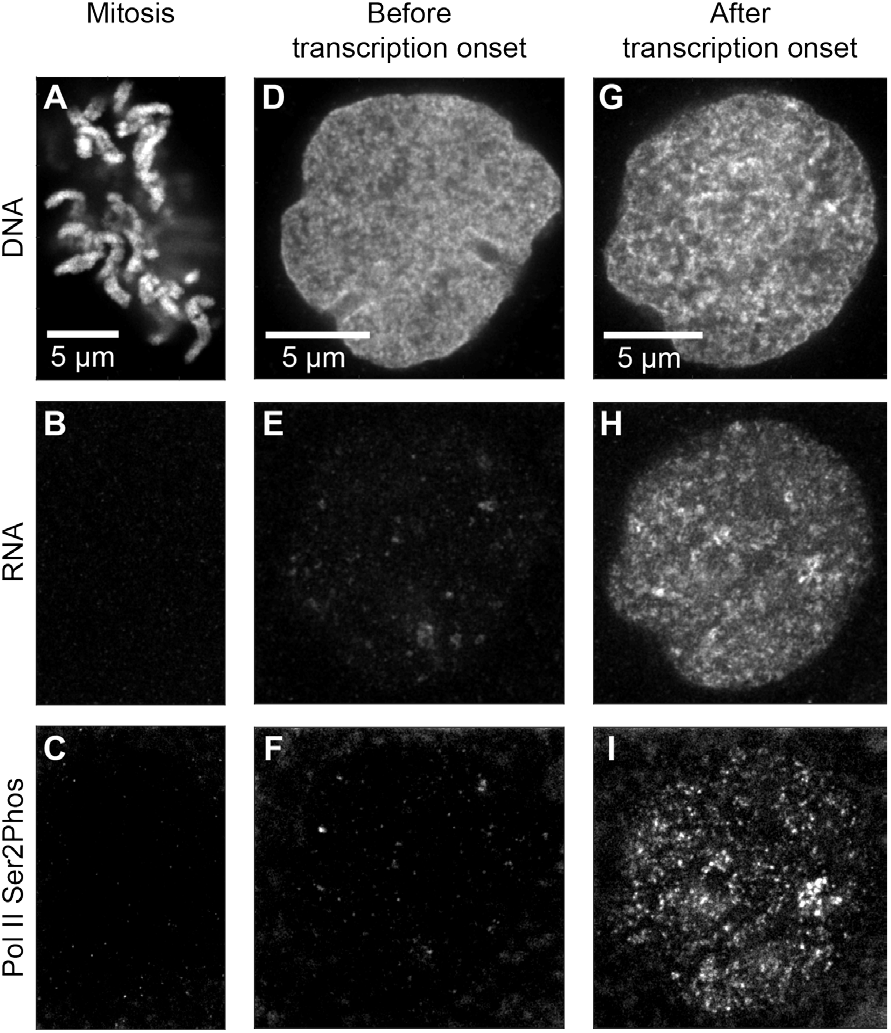
STED super-resolution micrographs of mitotic chromosomes, as well as nuclei before and after transcription onset. Representative three-color STED superresolution micrographs showing DNA, RNA, and transcriptional activity (Pol II Ser2Phos) during mitosis (A-C), shortly after mitosis (E-F), and in interphase after full transcription onset (G-I). All nuclei recorded from the same sample; acquisition settings and intensity maps are kept the same across all imaged nuclei.

**SI Figure 4.**
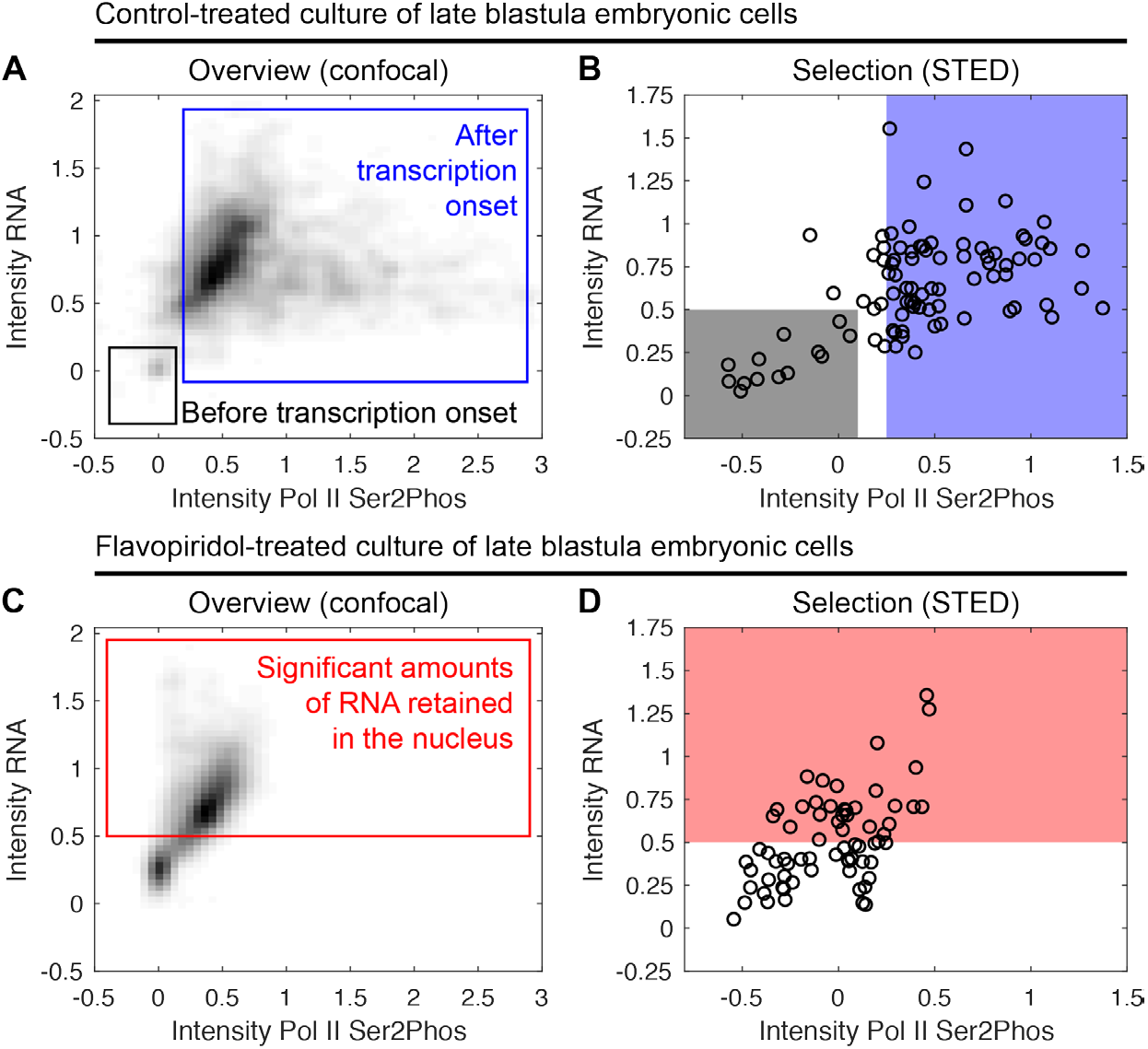
Selection of control-treated and flavopiridol-treated cells for image analysis. **A)** Empirical probability distribution of late blastula cells cultured in control media. Nuclear intensity of Pol II Ser2Phos and RNA were quantified from nuclear mid-sections acquired by spinning disk confocal microscopy in fixed cells. Cells before and after transcription onset are indicated by windows. Mitotic, prophase, and prometaphase nuclei were excluded by nuclear morphology and positive staining against Histone 3 Serine 28 phosphorylation. Data points pooled from 3 samples, intensities are scaled so that 0 and 1 correspond to the 5 and 95 percentiles of each sample. **B)** Plot representing individual nuclear mid-sections acquired by STED super-resolution microscopy, with selection windows corresponding to those in panel A. Data are pooled from 4 independent samples processed in two experiments, intensities scaled to the 5 and 95 percentiles of each experiment. Data from the two windows were used for Figure 1B, data from the “after transcription onset” window in Figure 1D,G and Figure 2E. **C)** Empirical probability distribution of cells cultured in flavopiridol containing media. Staining, acquisition, and analysis as in panel A. Cells retaining significant amounts of RNA in the nucleus are indicated by a window. Data points pooled from 3 samples, intensities are scaled to the 5 and 95 percentiles established from the control samples. All data points from this data set were used for Figure 2A. **D)** Plot representing individual nuclear mid-sections acquired by STED super-resolution microscopy, with selection window corresponding to that in panel C. Data are pooled from 4 independent samples processed in two experiments, intensities scaled to the 5 and 95 percentiles of the control samples of each experiment. Data from the window were used in Figure 2C,E.

**SI Figure 5.**
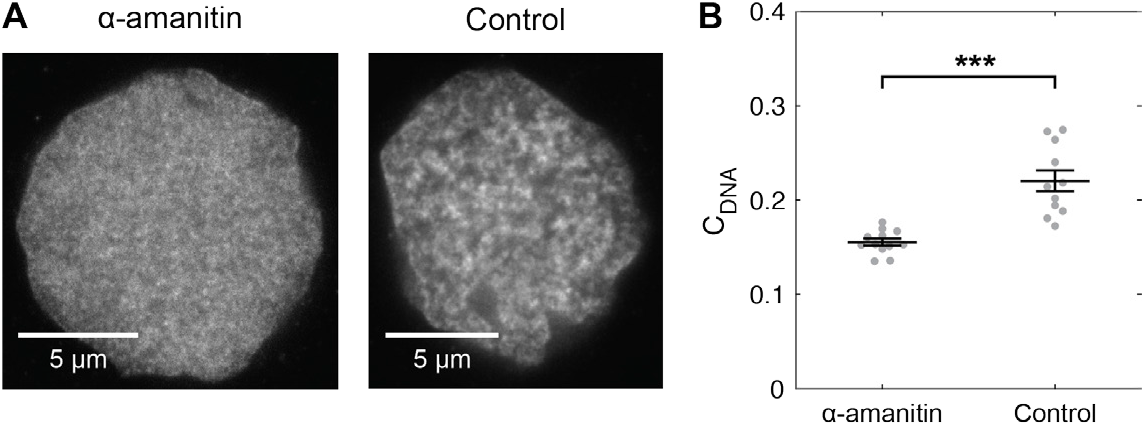
Transcription inhibition with α-amanitin suppresses the formation of DNA domains. **A)** STED super-resolution micrographs showing DNA intensities in interphase nuclear mid-sections of nuclei from α-amanitin injected and water-injected late blastula zebrafish embryos. **B)** DNA image contrast inside nuclei (C_DNA_), individual *C_DNA_* values with mean±SD; *** p<0.001 for difference of means, Bonferroni-corrected permutation test, n = 12, 11. Mitotic, prophase, and prometaphase nuclei were excluded by nuclear morphology and positive staining for Histone 3 Serine 28 phosphorylation.

**SI Figure 6.**
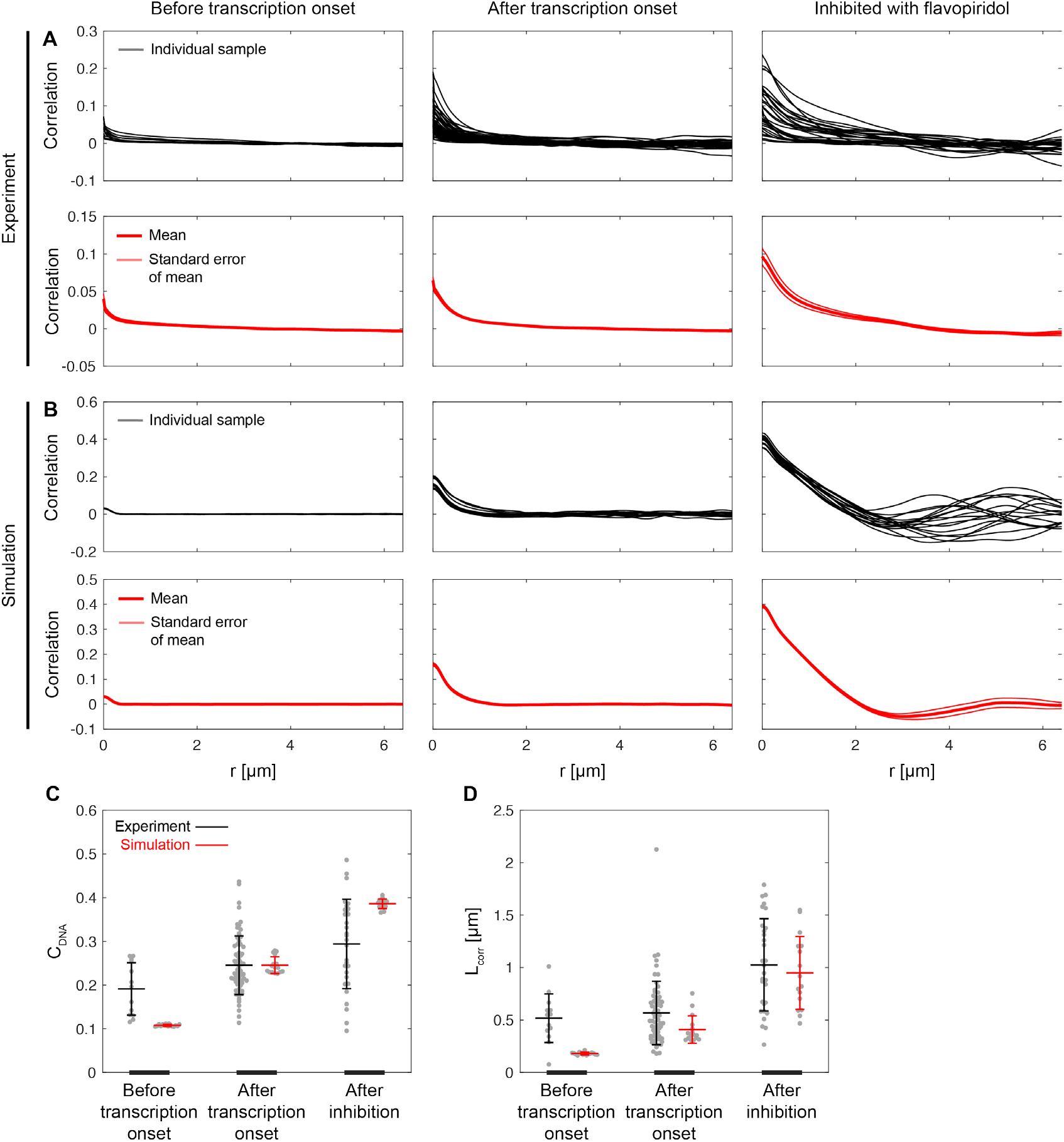
Quantification of spatial organization from microscopy DNA intensity profiles and simulated chromatin density profiles. **A)** Radial correlation functions of the DNA intensity profiles in STED super-resolution nuclear mid-sections in the indicated cells. **B)** Radial correlation functions from simulated chromatin density profiles under the indicated conditions. **C)** Image contrast (C_DNA_) in microscopy DNA intensity profiles (mean±std.dev., n=13, 66, 30 individual nuclei) and simulated chromatin density profiles (n=16 simulations). D) Correlation length (L_corr_) from the same microscopy DNA intensity profiles and simulated chromatin density profiles as in C.

**SI Figure 7.**
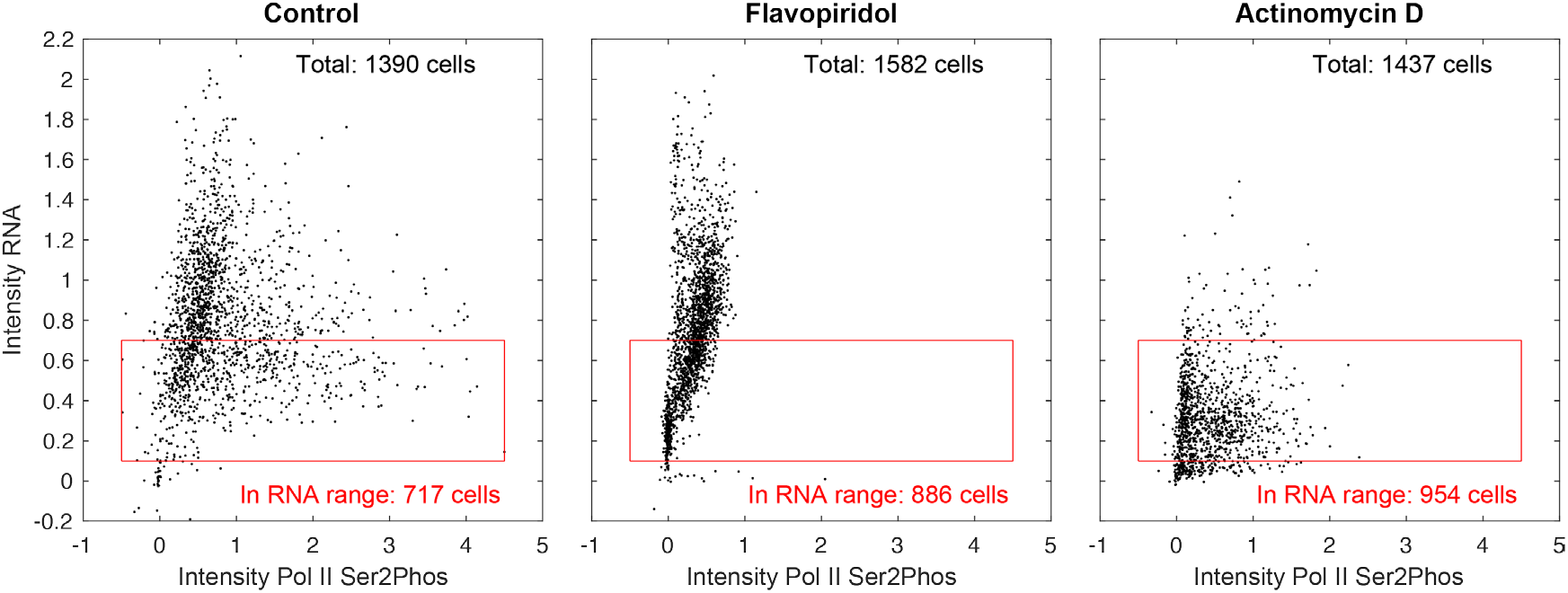
Effect of transcription inhibitors on transcriptional activity and RNA content in the nucleus. Dissociated cells were cultured in control media, media with flavopiriodol, and media with actinomycin D. Transcriptional activity (Pol II Ser2Phos) and RNA accumulation in the cell nucleus were quantified from fixed cells (data points pooled from 3 samples per condition, scaled so that 0 and 1 correspond to the 5 and 95 percentiles in the control sample). To allow a comparison of image contrast between the three different conditions while minimizing the influence of the RNA amount in the nucleus, a range of RNA intensities was defined that was present for all inhibitors. This range is indicated by the red frames, and only nuclear mid sections within these boxes are used for the comparison of image contrast in Figure 2F. Note that the different effects of flavopiridol and actinomycin D treatment can be expected based on their mechanisms of action. Flavopiridol prevents the transition of RNA polymerase II from initiation to elongation, but not elongation itself. Thus, elongating RNA polymerase II is lost as the transcription of genes is completed and no new elongation is established. RNA production, however, continues for a considerable length of time, while the elongation of currently transcribed genes is completed. Actinomycin D arrests elongating RNA polymerase II, which then remains bound to DNA and retains the Ser2Phos mark. Thus, inhibition by actinomycin D immediately stops further production of RNA transcripts but only a modest reduction in the Pol II Ser2Phos signal is seen.

**SI Figure 8.**
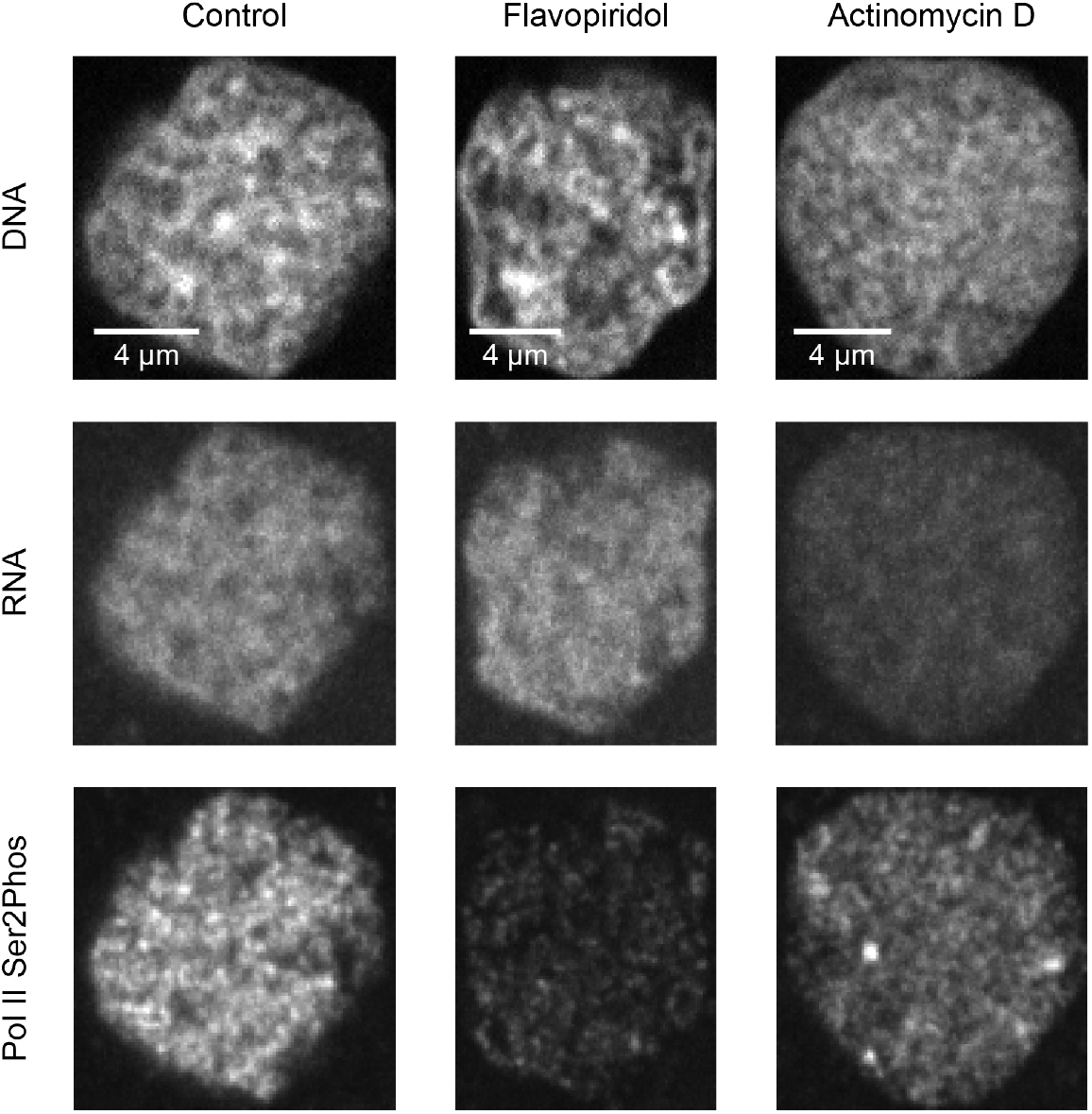
Actinomycin D treatment suppresses transcriptional activity, but a speckled pattern of transcriptional activity is retained. Example nuclear mid-sections from fixed cells after culturing in control media, media with flavopiridol, or media with actinomycin D. Micrographs recorded by spinning disk confocal microscopy. The different effects of flavopiridol and actinomycin D are expected. Flavopiridol prevents the transition of RNA polymerase II from initiation to elongation, but not elongation itself, so that currently transcribing RNA polymerase II (detected by the Pol II Ser2Phos mark) is only gradually lost as transcripts are completed. In consequence, RNA production is not rapidly interrupted. In contrast, actinomycin D rapidly arrests RNA transcript production, seen by the reduction of RNA in the nucleus, but the arrested RNA polymerase II presents as a pattern of Pol II Ser2Phos speckles.

**SI Figure 9.**
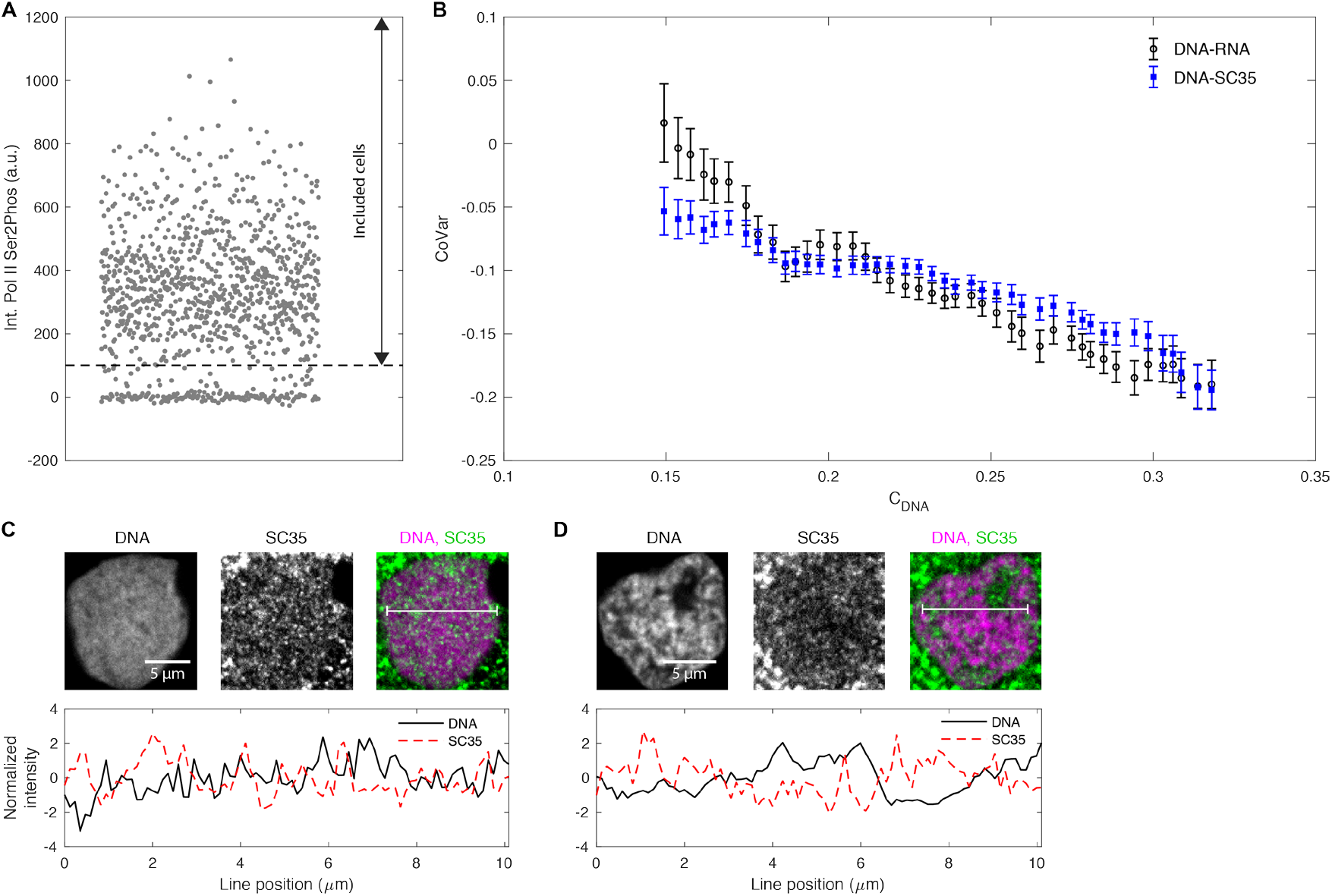
Formation of chromatin domains is accompanied by demixing of DNA and SC35. **A)** We tested whether the RNA binding protein SC35 and DNA demix in flavopiridol-treated cells. To obtain a more homogeneous cell population, we excluded cells without residual transcription activity (Pol II Ser2Phos), which have likely undergone mitosis after flavopiridol was applied, and therefore contain no nuclear RNA. All data were recorded by spinning disk confocal microscopy of fixed cells. Because no differing conditions are being compared, the arbitrary units here are taken directly from the intensity counts obtained during acquisition. Points are scattered in horizontal direction for visibility. **B)** The covariance (CoVar) between DNA and RNA as well as between DNA and the canonical splicing protein SC35 is shown for cells binned by increasing DNA image contrast in the nucleus (C_DNA_) in nuclear mid-sections (mean±s.e.m.). **C)** Example nuclear mid-section showing DNA and SC35 intensity profiles in a nucleus with low C_DNA_, and a color merge of both profiles. A line profile of both channels is also given. **D)** Example nuclear mid-sections of a nucleus with high C_DNA_, with line profile.

**SI Figure 10.**
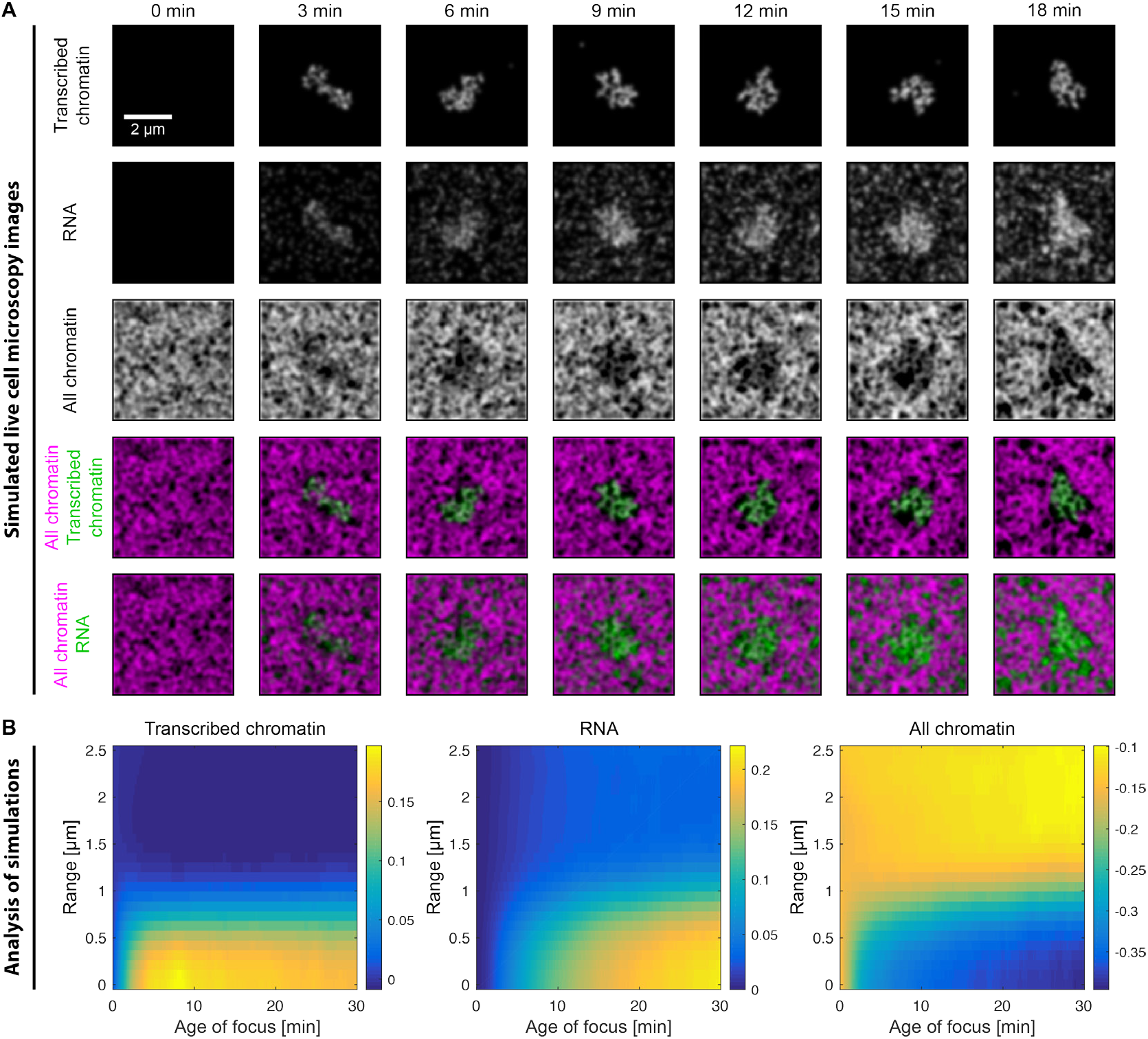
Dynamics of microenvironment formation in simulations of transcription onset at an isolated transcription site. The transcription onset at an isolated transcription site in a background of not transcribed chromatin is simulated using a 5-by-5 μm^2^ lattice (100 times 100 lattice sites). **A)** Representative time course of the different concentration profiles extracted from a single simulation. **B)** Radial analysis over 30 simulations. The range is relative to the centroid of transcribed chromatin, and extends outward in a radial fashion from that centroid. The age of the focus is counted from the point when the focus is first detected. Concentration values are background-subtracted, leading to negative values in some cases.

**SI Figure 11.**
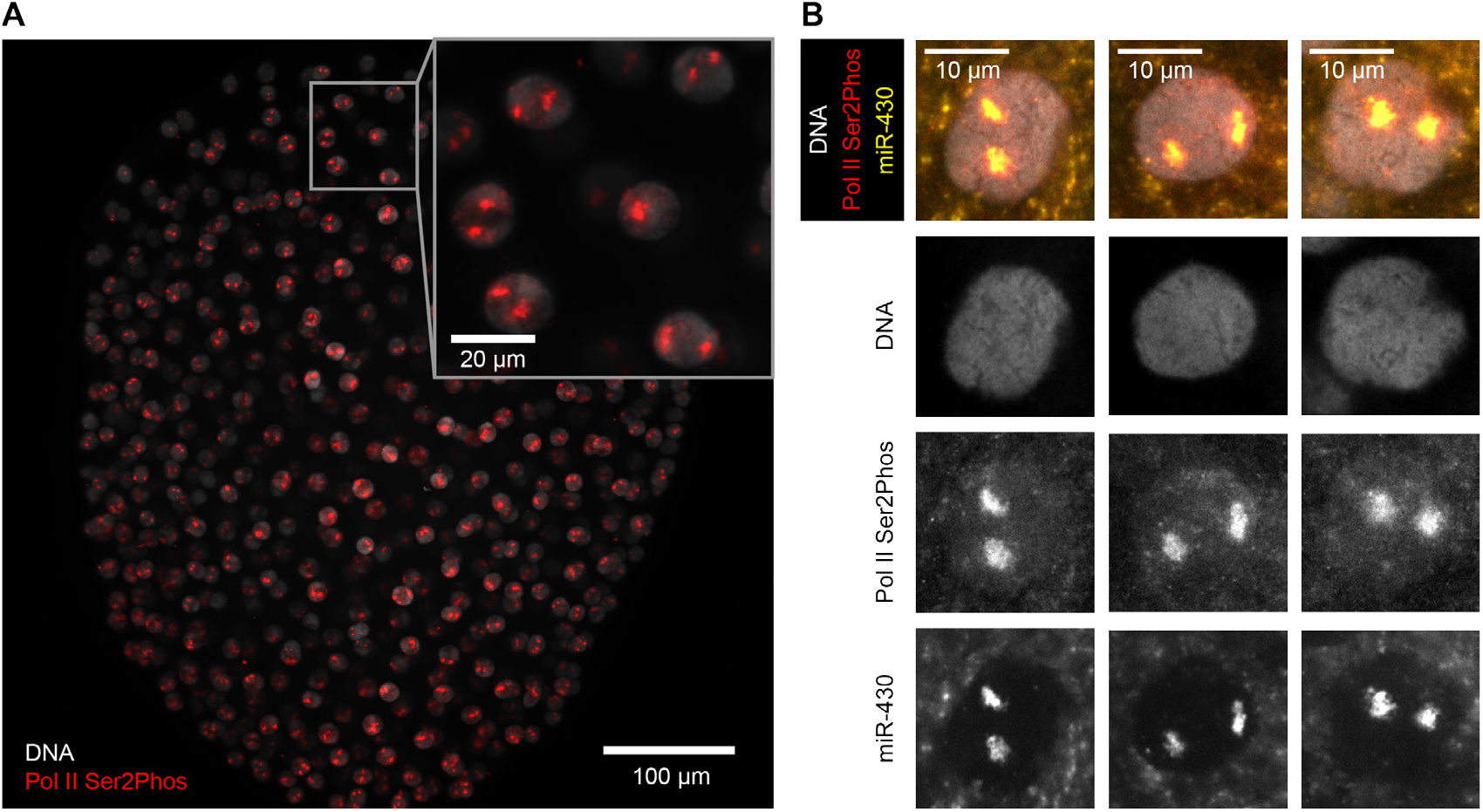
Two prominent transcription foci associated with the miR-430 gene cluster occur throughout the nuclei of late blastula zebrafish embryos. **A)** Light sheet micrograph showing the occurrence of two prominent transcription sites (Pol II Ser2Phos) in the nuclei (DNA labelled with DAPI) of a fixed late blastula zebrafish embryo (animal view, maximum intensity projection). **B)** Three example micrographs showing the colocalization of the two prominent transcription sites with miR-430 primary transcripts (pri-miRNA, revealed by fluorescence *in situ* hybridization, FISH) (van Boxtel *et al.*, 2015). Images acquired by spinning disk confocal microscopy, shown are single, mid-nuclear optical sections of three representative nuclei. Note that the FISH procedure can perturb the fine structure of chromatin, so that the depletion of DNA at the transcription sites is not as obvious as in images shown in other figures.

**SI Figure 12.**
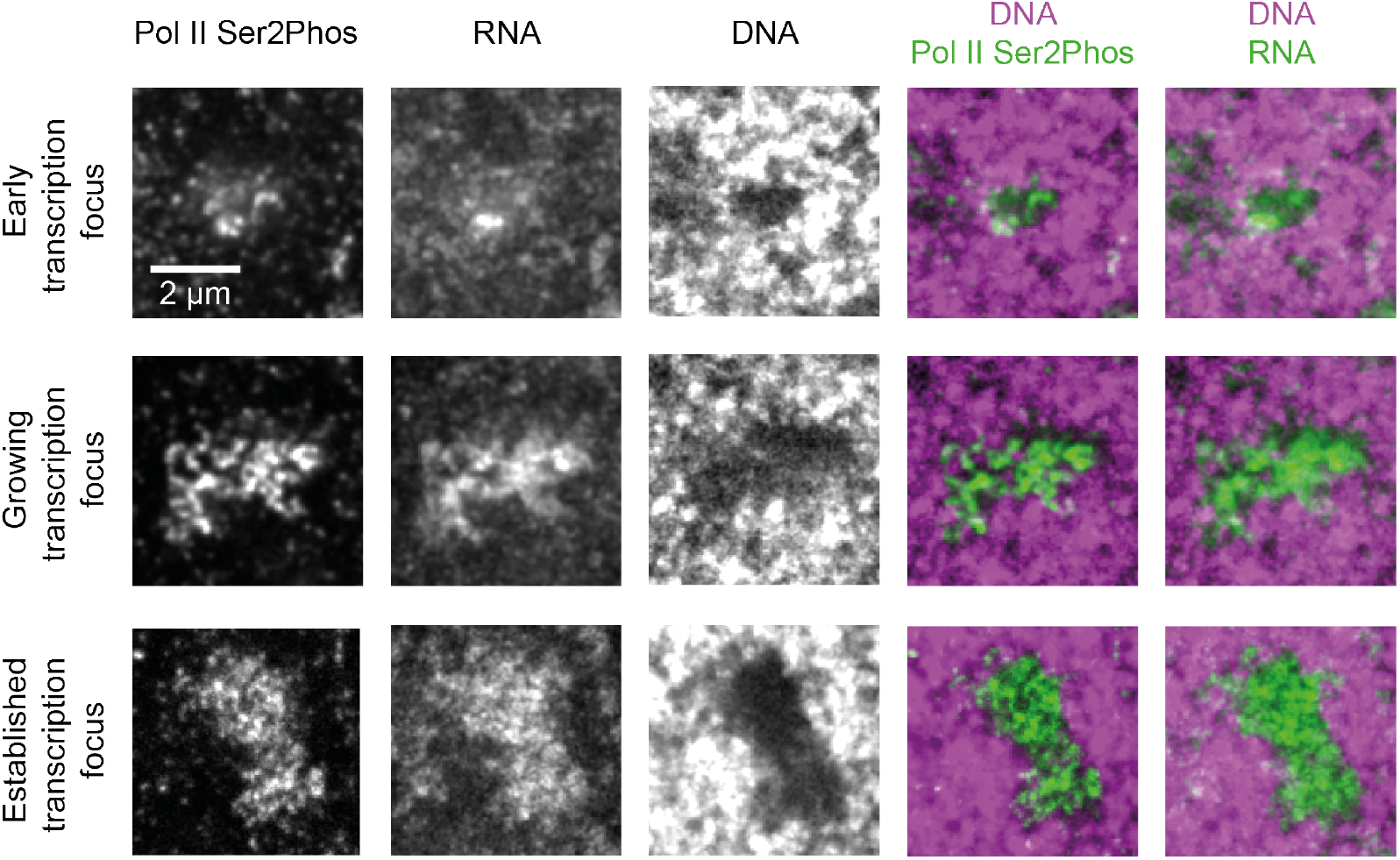
Super-resolution assessment of the establishment of prominent transcription foci. Representative zooms into STED super-resolution mid-sections of nuclei of fixed cells, showing prominent transcription in different stages of their emergence.

